# Site-specific lipidation enhances IFITM3 membrane interactions and antiviral activity

**DOI:** 10.1101/2020.09.11.293324

**Authors:** Emma Garst, Hwayoung Lee, Tandrila Das, Shibani Bhattacharya, Avital Percher, Rafal Wiewiora, Isaac P. Witte, Yumeng Li, Michael Goger, Tao Peng, Wonpil Im, Howard C. Hang

## Abstract

Interferon-induced transmembrane proteins (IFITMs) are *S*-palmitoylated proteins in vertebrates that restrict a diverse range of viruses. *S*-palmitoylated IFITM3 in particular directly engages incoming virus particles, prevents their cytoplasmic entry, and accelerates their lysosomal clearance by host cells. However, the precise molecular mechanisms of action for IFITM-mediated viral restriction are still unclear. To investigate how site-specific *S*-palmitoylation controls IFITM3 antiviral activity, here we employed computational, chemical, and biophysical approaches to demonstrate that site-specific lipidation of IFITM3 at highly conserved cysteine 72 modulates its conformation and interaction with lipid membranes leading to enhanced antiviral activity of IFITM3 in mammalian cells. Collectively, our results demonstrate that site-specific *S*-palmitoylation of IFITM3 directly alters its biophysical properties and activity in cells to prevent virus infection.

## INTRODUCTION

Interferon-induced transmembrane proteins (IFITMs) are *S*-palmitoylated proteins implicated in the immune response to viral infections (**Figure 1**). IFITMs were identified as interferon (IFN)-induced genes more than 30 years ago (Friedman, Manly, McMahon, Kerr, & Stark, 1984), but the broad antiviral activity of IFITM1, IFITM2, and IFITM3 was discovered more recently from siRNA knockdown (Brass et al., 2009) and overexpression screens (Schoggins et al., 2011). IFITM3 is the most active isoform and restricts many human pathogens entering through the endocytic pathway including influenza A virus (IAV) (Everitt et al., 2012), dengue virus (DENV) (Brass et al., 2009; John et al., 2013), Ebola virus (EBOV) (Brass et al., 2009; I.-C. Huang et al., 2011), Zika virus (Savidis et al., 2016), and other virus infections in cell culture (Bailey, Zhong, Huang, & Farzan, 2014; Perreira, Chin, Feeley, & Brass, 2013). IFITM3 is up-regulated by interferon stimulation in many cell types including cardiac fibroblasts (Kenney et al., 2019) and dendritic cells (Infusini et al., 2015), but is also constitutively expressed in most cell types, including the lung epithelium (X. Sun et al., 2016), embryonic stem cells (Xianfang Wu et al., 2018), and some tissue resident T cells (Wakim, Gupta, Mintern, & Villadangos, 2013; Wakim et al., 2012). Thus, IFITM3 provides both intrinsic and inducible protection against a variety of viral pathogens in many tissues. Beyond cell-intrinsic immunity, IFITMs have also been suggested to modulate adaptive immune responses through protection of immune effector cells from viral infection (Wakim et al., 2013) and by regulating CD4^+^ T-cell differentiation (Yánez et al., 2018). IFITMs reduce the susceptibility of trophoblasts to viral infection in the placenta while preventing trophoblast cell-cell fusion mediated by the ancestral retrovirus-derived syncytin protein, an essential process for fetal development (Buchrieser et al., 2019; Zani et al., 2019). Notably, infection of *Ifitm3-/-* mice with H3N2 IAV and pandemic H1N1 IAV led to increased morbidity and mortality (Bailey, Huang, Kam, & Farzan, 2012; Everitt et al., 2012; Kenney et al., 2019). IFITMs have also been shown to be expressed in other vertebrates and implicated in their resistance to viral infections (Bailey et al., 2014; Benfield et al., 2020; Compton et al., 2016). IFITMs are therefore clearly important for host susceptibility to diverse virus infections, which warrants further investigation into the mechanistic underpinnings of their antiviral activity.

**Figure 1.**
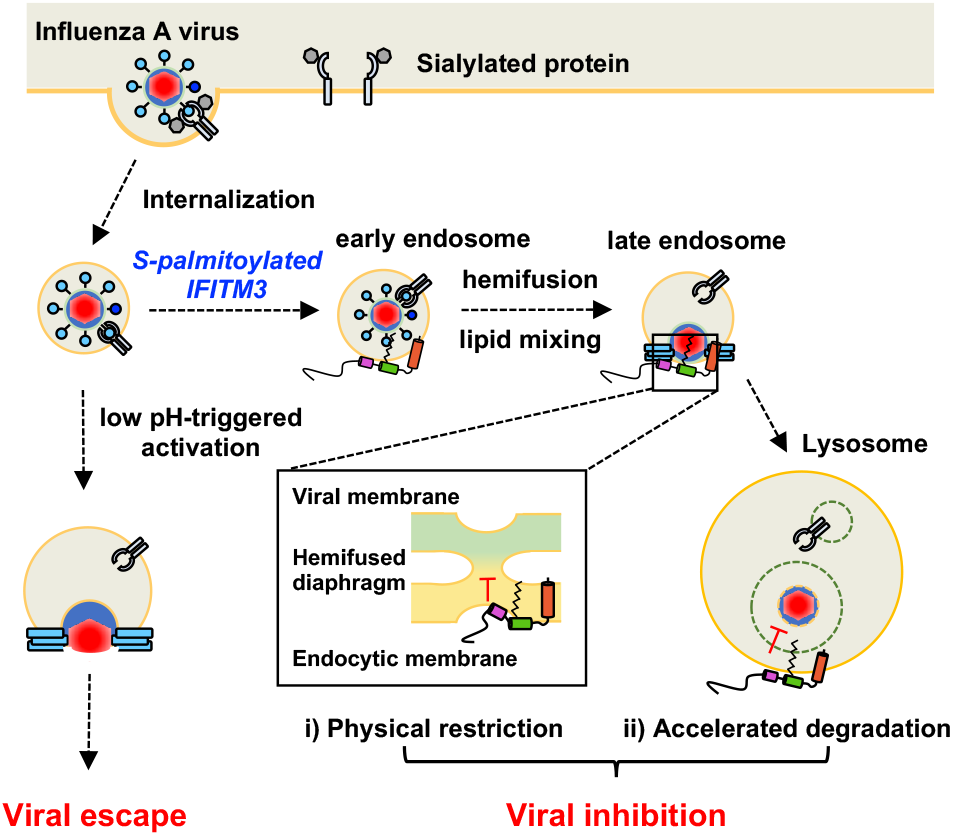
*S*-palmitoylation is essential for the antiviral activity of IFITM3. *S*-palmitoylated IFITM3 restricts viral entry through the endocytic pathway. In brief, a virus can enter the cell by binding a surface receptor and being internalized through the endocytic pathway. Without the presence of IFITM3, the virus can fuse with the endocytic membrane, releasing its contents into the cytoplasm for replication. When IFITM3 is *S*-palmitoylated at Cys72, it colocalizes with virus-containing endosomes in the early endocytic pathway and prevents the release of the virus into the cytoplasm (Spence et al., 2019; Suddala et al., 2019). Two non-exclusive models have been proposed for this activity: *i*) IFITM3 physically restricts the virus in the endocytic compartment preventing membrane fusion, perhaps by stabilizing a hemifused membrane intermediate, or *ii*) IFITM3 prevents the egress of the virus from the endocytic pathway by rapidly shuttling the endosome and its cargo for lysosomal degradation. The following figure supplement is available for figure 1: **Figure supplement 1**. Structure and lipidation of IFITM3.

In humans, a naturally occurring single nucleotide polymorphism (SNP rs12252) proposed to express an N-terminal truncated isoform of IFITM3 has been correlated with severe influenza in hospitalized populations (Everitt et al., 2012; Y.-H. Zhang et al., 2013); however, transcription of this isoform has not been detected by RNA sequencing of patient samples (Makvandi-Nejad et al., 2017; Randolph et al., 2017). Another loss-of-function allele has been identified in the IFITM3 gene 5’ untranslated region (SNP rs34481144), and is associated with lower IFITM3 mRNA expression and a decrease in airway resident CD8^+^ T cells during viral infection (Allen et al., 2017). Significantly, a recent small human cohort study showed that the rs12252 SNP was also associated with a higher incidence of severe COVID-19 (Y. Zhang et al., 2020b). Previously, overexpression of IFITM3 was shown to inhibit SARS-CoV infections (I.-C. Huang et al., 2011), but surprisingly promoted the infection of lung epithelial cells by the endemic human coronavirus OC43 (Zhao et al., 2014; 2018). Early studies on the role of the IFITMs in SARS-CoV-2 infection have been inconclusive. In initial screening and overexpression studies, IFITM2 and IFITM3 were found to inhibit SARS-CoV-2 pseudotyped virus (X. Zhang et al., 2020a), and IFITMs were shown to inhibit SARS-CoV-2 spike protein mediated cell-cell fusion (Buchrieser et al., 2020). However, conflicting studies have observed that SARS-CoV-2 pseudotyped virus infection (Zheng et al., 2020) or spike protein mediated cell-cell fusion (Zang et al., 2020) are not inhibited by IFITM3. More recent studies with genuine SARS-CoV-2 point to a more nuanced role in viral restriction. In these studies, IFITM overexpression was shown to restrict SARS-CoV-2 infection, though expression of an IFITM3 endocytosis mutant that accumulates at the plasma membrane enhanced SARS-CoV-2 infection (Shi et al., 2020). These results suggest that IFITM3 may have opposing effects on SARS-CoV-2 depending on the cellular location at which the virus fusion process occurs. Interestingly, IFITMs may have a proviral effect on SARS-CoV-2 infection of lung epithelial cells (Prelli Bozzo et al., 2020), which are infected primarily via plasma membrane fusion as opposed to endosomal fusion (Hoffmann, Kleine-Weber, et al., 2020a; Hoffmann, Mösbauer, et al., 2020b). As this potentially proviral effect is unique to coronaviruses, a mechanistic understanding of the activity of IFITMs will be key in delineating their roles in coronavirus infections.

Cellular studies of IFITMs have begun to reveal key features of their antiviral activity. Immunofluorescence (Chesarino, McMichael, & Yount, 2014a; Perreira et al., 2013) and live cell imaging (Peng & Hang, 2016) have shown that IFITM3 is largely localized to endolysomal vesicles in mammalian cells, where it restricts viruses that enter the cell through the endocytic pathway (Desai et al., 2014). Live cell imaging studies during virus entry by our laboratory (Spence et al., 2019) and others (Suddala et al., 2019) revealed that IFITM3 directly engages incoming virus-containing vesicles and accelerated their trafficking to lysosomes for destruction. Lipid mixing assays showed that IFITM3 does not prevent viral hemifusion with host membranes (Desai et al., 2014), suggesting that IFITMs may inhibit virus pore formation. Although the antiviral mechanism of IFITM3 is still unclear, two non-exclusive models for the restriction of viral particles have emerged (**Figure 1**). IFITM3 could actively accelerate the degradation of viral particles by manipulation of cellular trafficking pathways, as observed by live cell imaging studies (Spence et al., 2019). Alternatively, IFITM3 may physically restrict the virus infection by altering the biophysical properties of host membranes. This is supported by recent studies indicating that IFITM3 can increase the order and rigidity of endosomal membranes as well as induce negative curvature that could potentially stabilize a hemifused state while inhibiting viral pore formation (Guo et al., 2020; Rahman et al., 2020). Both of these mechanisms may account for the observed antiviral activity and cellular features of IFITM-expressing cells, including the enlargement and accumulation of cholesterol in endocytic vesicles (Amini-Bavil-Olyaee et al., 2013; Desai et al., 2014). Additional cellular and biophysical studies of IFITMs and their regulatory mechanisms are therefore still needed to understand their mechanisms of action.

IFITMs were originally proposed to be dual-pass transmembrane proteins (Brass et al., 2009), but epitope mapping studies in mammalian cells (Bailey, Kondur, Huang, & Farzan, 2013; Weston et al., 2014; Yount, Karssemeijer, & Hang, 2012) and *in vitro* electron paramagnetic resonance (EPR) spectroscopy and nuclear magnetic resonance (NMR) spectroscopy studies (Ling et al., 2016) have indicated that IFITM3 is a type IV single-pass transmembrane protein with an amphipathic region from residue Trp60 to Arg85 containing two α-helices (**Figure 1 – figure supplement 1**). This region is contained within the conserved CD225 domain, which also includes an intrahelical loop from residue Arg85 to Lys104 (El-Gebali et al., 2018). Through alanine scanning mutagenesis, a number of point mutations within these regions at post-translational modification (PTM) sites or within putative oligomerization motifs were found to disrupt the antiviral activity of IFITM3 (John et al., 2013). Moreover, mutations within the first amphipathic helix (Val59-Met68) that altered IFITM3 hydrophilicity abrogated the antiviral activity, indicating that the amphipathic nature of this helix is required for its antiviral function (Chesarino et al., 2017). Furthermore, amphipathic helix 1 can alter the biophysical characteristics of the phospholipid bilayer by inducing negative curvature, increasing lipid order, and increasing membrane stiffness (Guo et al., 2020). These studies confirm that the conserved amphipathic region of IFITM3 is necessary for the restriction of viral particles in the endocytic pathway, possibly through the direct manipulation of the local membrane environment.

IFITM3 activity is regulated by a number of PTMs such as *S*-palmitoylation (Cys71, Cys72, and Cys105), ubiquitination (Lys24, Lys83, Lys88, and Lys104), phosphorylation (Tyr20), and methylation (Lys88) (Chesarino et al., 2014a). The interplay between these PTMs has been implicated in the localization, activity, and turnover of IFITM3. Methylation and ubiquitination at Lys88 can both down-regulate IFITM3 activity (Yount et al., 2012), whereas phosphorylation at Tyr20 can block IFITM3 endocytosis and ubiquitination (Chesarino, McMichael, Hach, & Yount, 2014b). Ubiquitination at Lys24 promotes the interaction of IFITM3 and VCP/p97, which regulates IFITM3 trafficking and turnover (Xiaojun Wu et al., 2020). Several studies from our laboratory have demonstrated IFITM3 can be *S*-palmitoylated at Cys71, Cys72, and Cys105 (Percher et al., 2016; Yount et al., 2012; 2010). In particular, Cys72 is highly conserved across most mammals and is required for the antiviral activity of IFITM3 orthologs from mice, bats, and humans (Benfield et al., 2020; John et al., 2013; Percher et al., 2016). Site-directed mutagenesis and live cell imaging studies by our laboratory and others have shown that residue Cys72 is essential for IFITM3 antiviral activity, trafficking, and colocalization with incoming viral particles in the endocytic pathway (Spence et al., 2019; Suddala et al., 2019). Moreover, overexpression of S-palmitoyltransferases (zDHHC-PATs 3, 7, 15, and 20) led to increased IFITM3 *S*-palmitoylation and antiviral activity (McMichael et al., 2017). These studies demonstrate that site-specific and regulated *S*-palmitoylation of IFITMs is particularly crucial for their antiviral activity, but these studies were primarily from loss-of-function phenotypes and did not demonstrate if site-specific lipidation confers gain-of-function.

*S*-Palmitoylation is a reversible PTM in eukaryotes. *S*-Palmitoylation targets peripheral membrane proteins to specific cellular membranes (Rocks et al., 2010) and can act as a sorting mechanism in cells without specific receptor-ligand pairing (Rocks et al., 2005). For transmembrane proteins, *S*-palmitoylation can promote or disrupt association with specific membrane microdomains (Abrami, Leppla, & van der Goot, 2006; Levental, Lingwood, Grzybek, Coskun, & Simons, 2010; Yang et al., 2004), stabilize or disrupt protein-protein interactions (Lakkaraju et al., 2012; Yang et al., 2004; Zevian, Winterwood, & Stipp, 2011), and change the conformation of proteins (Abrami, Kunz, Iacovache, & van der Goot, 2008). Although significant advances have been made to detect and discover *S*-palmitoylated proteins (Hannoush & Sun, 2010; Yount et al., 2010), the functional analysis of site-specific *S*-palmitoylation, which may be sub-stoichiometric in cells, is still challenging. To investigate how *S*-palmitoylation enhances the antiviral activity of IFITMs, we employed *in silico* as well as chemical strategies to evaluate site-specifically lipidated IFITM3 structure *in vitro* and antiviral activity in mammalian cells. Our molecular dynamics simulation studies, the reconstitution of chemically lipidated IFITM3 *in vitro* (**Figure 1 – figure supplement 1**), and solution-state NMR spectroscopy analysis suggest that site-specific *S*-palmitoylation of IFITM3 at Cys72 induces conformational changes in the amphipathic helices and N-terminal domain, and anchor these cytoplasmic domains to cellular membranes. To complement these *in silico* and *in vitro* structural studies, we used genetic code expansion and bioorthogonal ligation methods with tetrazine-lipid analogs that our laboratories recently developed (**Li Y et al in review, Supplemental File**) for site-specific lipidation of IFITM3 in mammalian cells (**Figure 1 – figure supplement 1**). These studies showed that site-specific chemical lipidation enhanced IFITM3 antiviral activity in mammalian cells, providing additional support for our modeling and structural studies *in vitro*. Collectively, our studies highlight the importance of site-specific lipidation methods to investigate gain-of-function phenotypes for *S*-palmitoylation and underscores the significance of this PTM for the antiviral activity and biochemical properties of IFITMs.

## RESULTS

### Molecular dynamics simulation of S-palmitoylated IFITM3 and chemically lipidated variants

To understand how site-specific *S*-palmitoylation controls the interaction of IFITM3 with the membrane bilayer, we used molecular dynamics to simulate IFITM3 in a variety of lipidation states. For these studies, human IFITM3 was modeled in a 1,2-dimyristoyl-*sn*-glycero-3-phosphocholine (DMPC) bilayer without palmitoylation (apo), with a palmitoyl group at Cys72 (Cys72-Palm), or with a stable maleimide-palmitoylation mimic (Cys72-mPalm) (**Figure 2A**). We found that S-palmitoylation at Cys72 altered the positioning of the amphipathic region of IFITM3, particularly within the first amphipathic helix (AH1, Leu62-Phe67) (**Figure 2A**). To quantify and compare the effects of palmitoylation, we calculated the distance of the center of mass of each helix from the membrane center (at Z = 0) throughout the 1-μs simulation (**Figure 2B**). When unmodified, AH1 remained water exposed with an average distance of 26.5 Å away from the membrane center. Conversely, Cys72 palmitoylation led to increased membrane proximity in AH1, reducing its average distance from the membrane center to 16.1 Å near the membrane head group. The Cys72-mPalm modification also brought AH1 into close proximity with the membrane bilayer, showing an average distance of 9.1 Å (below the head group). Although similar in effect, the closer association of the mPalm-modified construct suggests that the maleimide head group of the palmitoyl mimic may either be able to insert AH1 better into the membrane bilayer or form stronger interactions with the phospholipids compared to the thioester of natural *S*-palmitoylation. Interestingly, AH2 in all models showed similar orientation and positioning regardless of Cys72 palmitoylation despite its proximity to the lipid modification. Thus, *S*-palmitoylation may lead to localized and specific changes in protein structure and dynamics.

**Figure 2.**
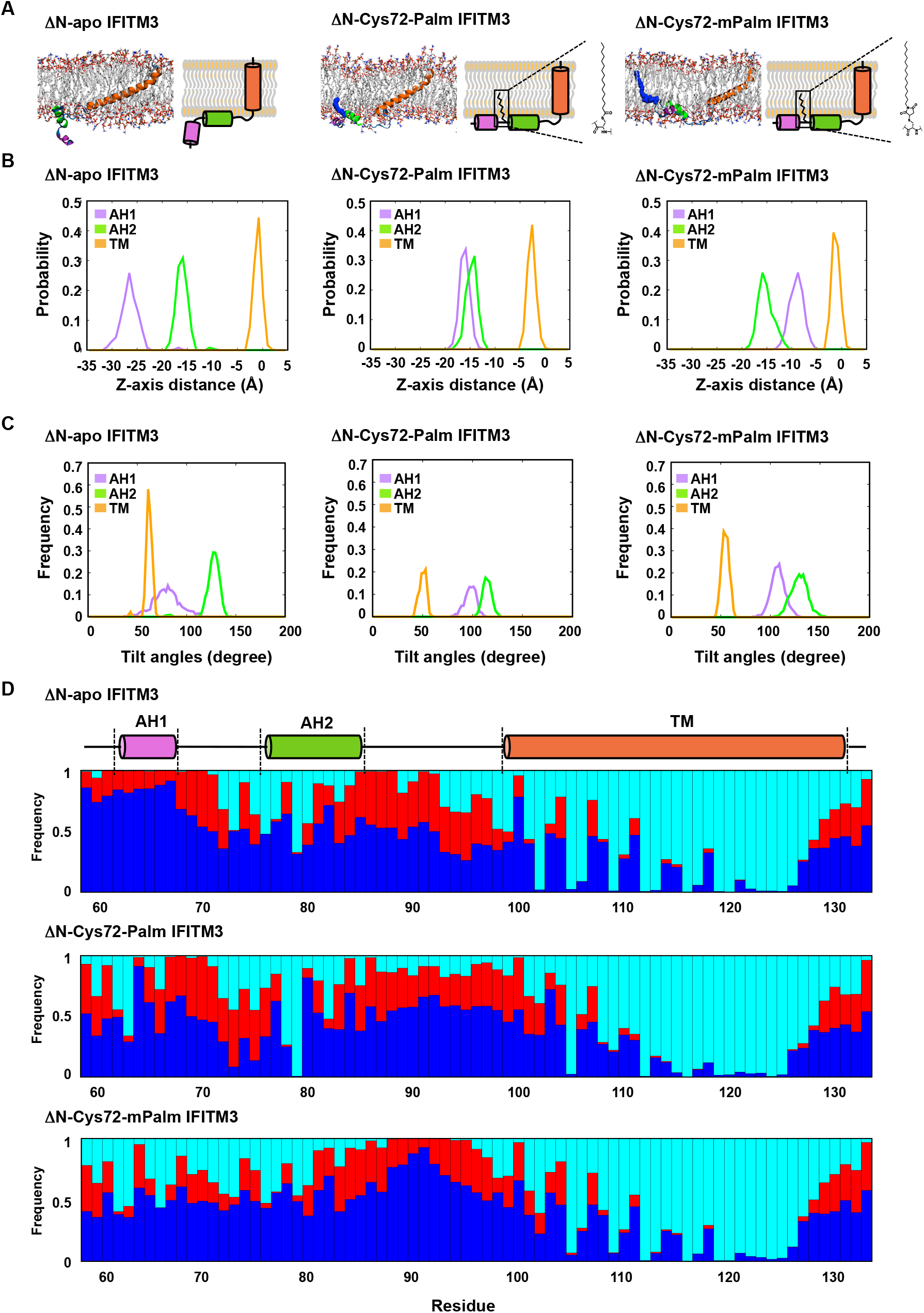
Molecular dynamics simulation of site-specifically lipidated IFITM3. A) Modeling snapshots of apo, Cys72-*S*-palmitoylated (Cys72-Palm), and Cys72-maleimide-palmitate (Cys72-mPalm) modified IFITM3 from residue Val58 to Gly133 (denoted as ΔN in figure) in DMPC bilayers are shown side by side with representative cartoons. The initial secondary structure was modeled based on published structural data (Ling et al., 2016), with amphipathic helix 1 from residue Leu62 to Phe67 (AH1, magenta), amphipathic helix 2 from residue Phe78 to Ser84 (AH2, green), and a 33 amino acid transmembrane domain from residue Ala98 to Ala131 (TM, orange). Molecular dynamics simulations were performed on these conditions for 1 μs, of which the last 500 ns were used for analysis. The simulations were run in triplicate for better sampling. B) The distance of the center of mass for each helix from the membrane center (Z = 0) was measured throughout the course of the simulation. Probability plots of the AH1, AH2, and TM domain position relative to the membrane center for the different IFITM3 variants are shown. The center of mass distance of AH1 is represented by magenta, AH2 by green, and the TM helix is represented by orange. C) The frequency with which IFITM3 AH1, AH2, and TM helix sample a specific helical tilt in regard to the membrane normal (the Z axis) was measured throughout the course of a 1-µs simulation. The helical tilt frequency of AH1 is represented by magenta, AH2 by green, and the TM helix is represented by orange. D) The frequency with which each residue of apo IFITM3, Cys72-Palm IFITM3, and Cys72-mPalm IFITM3 interacted with different environments throughout the course of the molecular dynamics simulation was measured. The interaction pattern graph shows the probability of occurrence within 4 Å from water (blue), DMPC lipid head (red), or DMPC lipid tail (cyan).

In addition to AH1 membrane association, we also measured the helical tilt of the α-helical segments (with respect to the membrane normal) throughout the simulations. These data show that when IFITM3 is not palmitoylated, AH1 is a flexible segment that can sample many orientations (**Figure 2C**). Additionally, during the simulation, the AH1 region frequently contacted water molecules, while only occasionally contacting the phospholipid head groups, as might be expected of a soluble peptide rather than one with amphipathic tendencies (**Figure 2D**). In contrast, when IFITM3 is modified at Cys72 either with a naturally occurring palmitoyl group or maleimide-palmitate, the AH1 is restricted to the surface of the model membrane, sampling a tight range of helical tilts around 90° from the membrane normal and interacting with water, the phospholipid head groups, and the lipid tails in relatively equal parts (**Figure 2C, D**). In the case of loop 2 (residue 86 to 97), the apo model interacts much more frequently with the phospholipid head groups compared to the lipidated models (Cys72-Palm and Cys72-mPalm), indicating the effect of lipidation on IFITM3 dynamics can be seen on both the N- and C-terminal side of the modification up to 20 residues away (**Figure 2D**). No notable difference was observed in the interactions between the transmembrane domain and its surrounding environments throughout all systems. Together, these data indicate that lipidation at Cys72 may be required for the proper folding and anchoring of IFITM3 AH1 despite its relative distance from AH1, and more significantly its location past a short disordered region. Moreover, this molecular simulation indicates that maleimide-palmitate can function as a reasonable mimic of *S*-palmitoylation for our structural and biochemical studies.

### Generation of recombinant site-specifically lipidated IFITM3

To further understand how site-specific lipidation regulates IFITM3, we optimized the expression and purification of recombinant human His_6_-tagged IFITM3 from *E. coli* and employed maleimide-palmitate (mPalm) for site-specific lipidation of key Cys residues (**Figure 3A**). Maleimide-palmitate was used due to its cysteine specific reactivity and potential for full length protein labeling in detergents. In order to optimize the maleimide coupling reaction, we tested a number of conditions including time, temperature, maleimide-palmitate concentration, and buffer composition, and monitored protein labeling by mass shift on an SDS-PAGE gel (**Figure 3 – figure supplement 1**). Through these studies, we found that the maleimide coupling reaction went to completion in 2 hours at 15°C, with a final maleimide-palmitate concentration of 0.5 mM in 10% DMSO. The reaction could be effectively blocked or quenched with excess *N*-ethyl maleimide or β-mercaptoethanol, respectively, indicating that the modification is specific to free cysteines (**Figure 3 – figure supplement 2**). Using this approach, we utilized Cys to Ala mutagenesis to generate a panel of site-specifically lipidated His_6_-IFITM3 constructs modified at Cys72, Cys105, and dually at Cys72 and Cys105. The maleimide coupling reaction was specific and went to completion, as shown through SDS-PAGE gel shift and MALDI analysis (**Figure 3B, C**). Comparison between His_6_-apo IFITM3 and His_6_-Cys72-mPalm IFITM3 reveals a mass difference of 362 Da, indicating the addition of one maleimide-palmitate group (**Figure 3C** and **Figure 3 – figure supplement 3**). The dually modified His_6_-Cys72Cys105-mPalm had a mass increase of 710 Da, consistent with the addition of two maleimide-palmitate groups (**Figure 3C** and **Figure 3 – figure supplement 3**). Together, these data show that IFITM3 can be effectively modified with the *S*-palmitoylation analog maleimide-palmitate at free cysteines under mild aqueous conditions.

**Figure 3.**
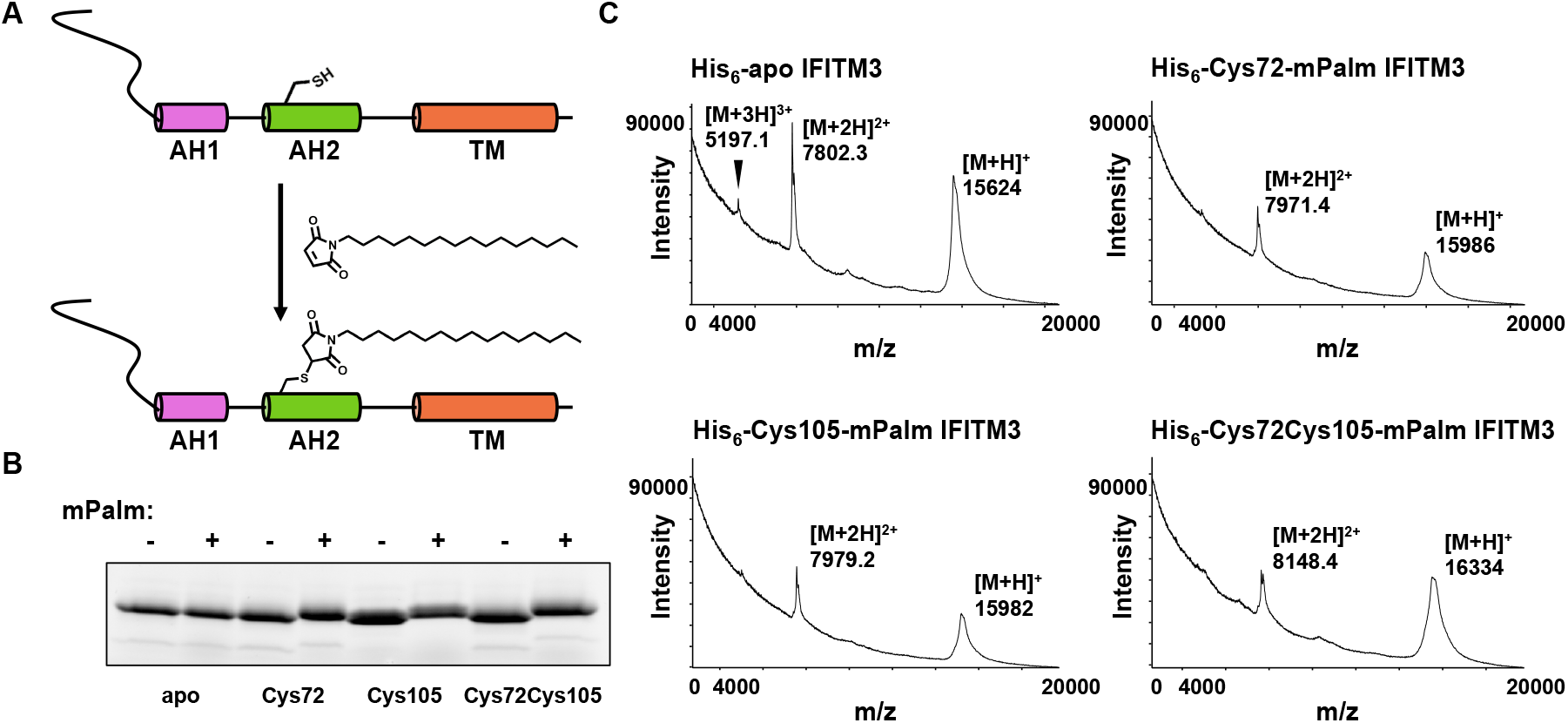
Generation and characterization of site-specifically lipidated IFITM3 *in vitro*. A) Schematic for the production of site-specifically lipidated IFITM3 via a maleimide coupling reaction specific for reduced cysteine residues. B) The successful coupling of maleimide-palmitate (mPalm) to IFITM3 at free cysteines was measured by an SDS-PAGE gel shift. The reaction went to completion in 2 hours at 15°C under mild reducing conditions (25mM HEPES pH 7.4, 150 mM KCl, 0.8% TX-100, 1mM TCEP, 0.5 mM maleimide-palmitate, 10% DMSO). C) MALDI spectra of apo and site-specifically mPalm-modified IFITM3 were collected in linear, delayed extraction mode with a delay of 0.75-1 μs and sampling rate of 2 ns. Each spectrum corresponds to approximately 1000 scans. Samples were calibrated internally with a horse myoglobin protein standard. The following figure supplements are available for figure 3: **Figure supplement 1**. Optimization of Cys72-IFITM3 maleimide-palmitate coupling reaction. **Figure supplement 2.** Maleimide-palmitate coupling with Cys72-IFITM3 can be blocked or quenched by *N*-ethyl maleimide (NEM) or β-mercaptoethanol (BME) respectively. **Figure supplement 3.** MALDI quantification of His6-apo IFITM3 and panel of His6-IFITM3 lipidated variants.

### Solution state NMR analysis of chemically lipidated IFITM3

To determine the effect of lipidation on the structure of His_6_-IFITM3, we expressed and purified isotopically labeled unmodified (apo) His_6_-IFITM3 and Cys72 monolipidated (Cys72-mPalm) His_6_-IFITM3 in dodecyl phosphocholine (DPC) micelles at pH7 for structural characterization by solution-state NMR (**Figure 4A**). In brief, isotopically labeled (^2^H/^13^C/^15^N) His_6_-apo and His_6_-Cys72-mPalm IFITM3 were purified and dialyzed into NMR buffer (25mM HEPES pH7, 150mM KCl, 0.5% DPC) for backbone assignments. A suite of ^15^N-traverse relaxation optimized spectroscopy (TROSY) based triple resonance experiments (^15^N-HSQC, HNCA, HNCO, HNCACB, HNCOCA, HNCOCACB, and HNCACO) were used to assign the backbone resonances. Sequential assignments were facilitated by ^15^N specific labeling of specific residues (Ala, Leu, Val, Ile, and Phe) to alleviate resonance overlap and allow the assignment of residues proximal to the site of lipidation (**Figure 4B** and **Figure 4 – figure supplement 1**). In total, we were able to assign 70% and 75% of the His_6_-Cys72-mPalm and His_6_-apo IFITM3 protein backbones respectively (**Figure 4 – figure supplement 2** and **Figure 4 – figure supplement 3)**. The backbone chemical shifts were analyzed in TALOS+ (Shen, Delaglio, Cornilescu, & Bax, 2009) to determine the secondary structure of His_6_-IFITM3, which was highly α-helical in the C-terminal domain (residue Ser61 to Gly133) (**Figure 4C**). In the N-terminal region, both His_6_-apo and His_6_-Cys72-mPalm IFITM3 had a high percentage of dynamic residues (26.7% and 35%, respectively). The random coil index (RCI) predicted order parameter S^2^ confirmed that the N-terminal domain of both protein constructs is highly dynamic (**Figure 4 – figure supplement 4**) (Berjanskii & Wishart, 2005).

**Figure 4.**
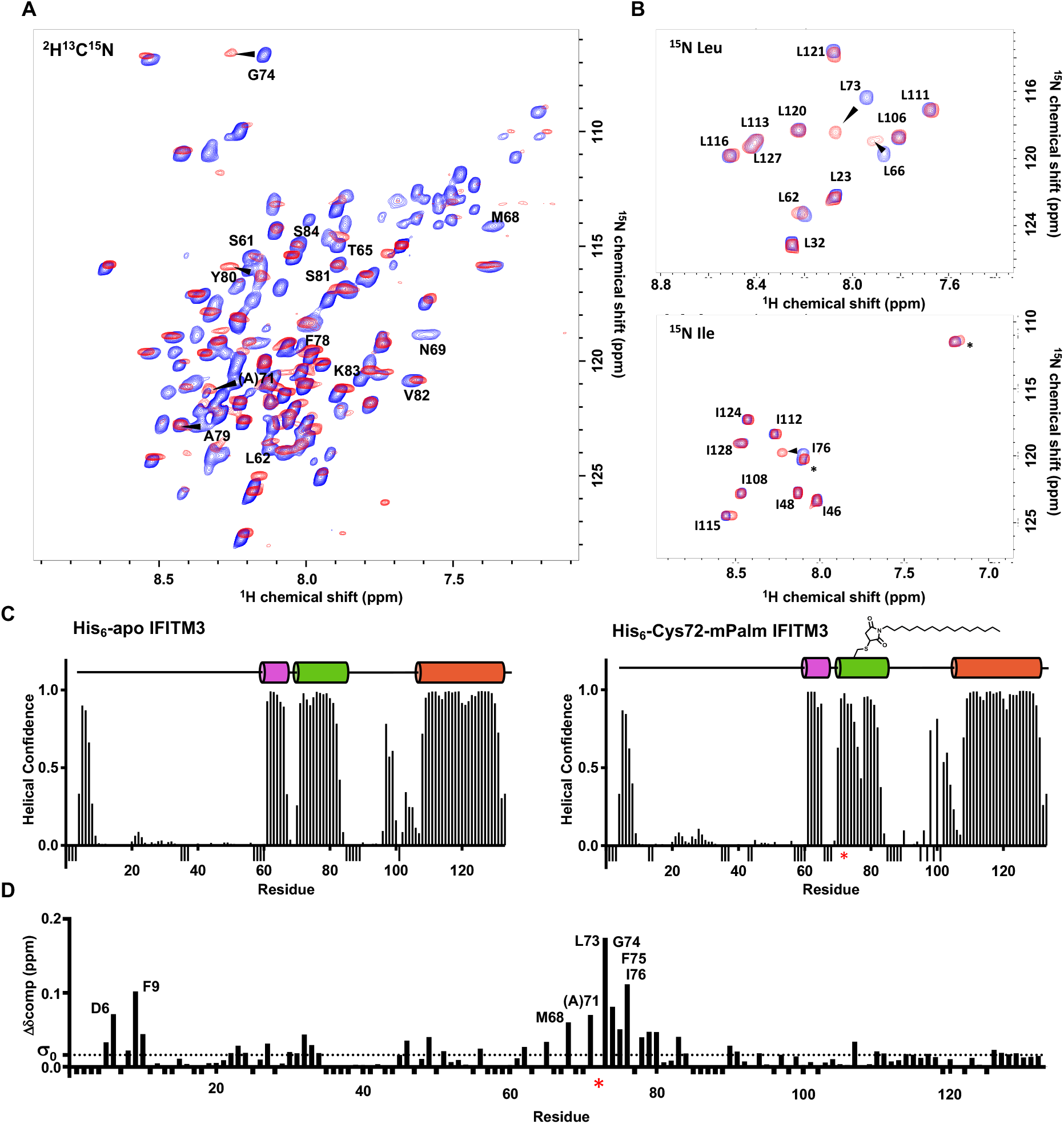
Comparative structural analysis of apo and C72-mPalm IFITM3 using NMR. A) ^1^H-^15^N TROSY spectra were recorded using ^2^H^13^N^15^C uniformly-labeled His_6_-apo (blue) and His_6_-Cys72-mPalm (red) IFITM3 in DPC micelles. Representative peaks from the conserved amphipathic region (Trp60 to Arg85) are labeled. B) ^1^H-^15^N TROSY spectra were collected of specifically labeled ^15^N-Leu and ^15^N-Ile His_6_-apo (blue) and His_6_-Cys72-mPalm (red) IFITM3. Arrows indicate changes in chemical shift (quantified in Figure 4D). Asterisks note residues without backbone assignment. C) The secondary structures of His_6_-apo and His_6_-Cys72-mPalm IFITM3 were predicted using TALOS+ based on chemical shift and torsional angle information. Residues without assignment due to low signal or spectral overlap are demarcated by negative bars to distinguish from residues without any helical propensity. The red asterisk indicates the site of mPalm modification. D) The magnitude of the change in chemical shift between modified and unmodified IFITM3 was measured by chemical shift perturbation analysis of His_6_-apo vs. His_6_-Cys72-mPalm IFITM3. Prolines and residues without assignment from either the His_6_-apo or His_6_-Cys72-mPalm IFITM3 structural analysis are demarcated by negative bars to distinguish from residues without any change in chemical shift. The standard deviation (σ_0_, dotted line) of the net change in chemical shift was calculated and used as a threshold value. Residues with a greater than 3σ change in chemical shift are labeled. The following figure supplements are available for figure 4: **Figure supplement 1**. ^15^N specific labeling of His_6_-apo and His_6_-Cys-mPalm IFITM3 for backbone assignment. **Figure supplement 2**. Backbone resonance assignment of His_6_-apo IFITM3 overlayed on ^1^H-^15^N TROSY spectra. **Figure supplement 3**. Backbone resonance assignment of His_6_-Cys72-mPalm IFITM3 overlayed on ^1^H-^15^N TROSY spectra. **Figure supplement 4**. Random coil index (RCI) analysis of His_6_-apo IFITM3 and His_6_-Cys72-mPalm IFITM3.

In the ordered C-terminal domain, our secondary structure analysis revealed three α-helices similar to previously published structural work on His_6_-IFITM3 (Ling et al., 2016). However, we found a number of notable differences from this published work in both the amphipathic region and the transmembrane domain. Two helices were observed in the amphipathic region for both the His_6_-apo-IFITM3 and His_6_-Cys72-mPalm-IFITM3 constructs: AH1 from Ser61 to Phe67 and AH2 from Cys71 to Lys83 (**Figure 4C**). In our analysis, AH2 was three residues longer than the published structure (which stretches from Ile76 to Arg85) and now contained the functional lipidation site, Cys72. Moreover, our data suggest a loop region from Lys104 to Asn107 followed by a 23-amino acid helix between Ile108 and Ala131 (**Figure 4C**), differing from the previously described 33-amino acid transmembrane domain (Ala98-Ala131) (Ling et al., 2016).

When comparing the secondary structure of our chemically lipidated and non-lipidated constructs, we found subtle changes in the amphipathic region while the global secondary structure remained the same. Lipidation at Cys72 caused a small decrease in the α-helical confidence directly C-terminal to the site of modification (Phe75 to Ala77) (**Figure 4C**). Furthermore, lipidation at Cys72 increased the helical propensity of residues Ala100 to Lys104, possibly indicating the stabilization of a third amphipathic helix in the region. These changes are indicative of a localized effect on the structure of IFITM3; however, lipidation does not destabilize the existing secondary structure of IFITM3 overall, consistent with our molecular dynamics simulations (**Figure 2**).

We also used chemical shift perturbation (CSP) analysis to quantify the effect of lipid modification on the structure of His_6_-IFITM3 (**Figure 4D**). This analysis utilizes the change in chemical shift of the amide backbone peaks to measure the magnitude of effect for some change to the protein or environment, in this case the addition of a lipid analogue at Cys72. This analysis revealed that maleimide-palmitate modification at Cys72 had a significant effect on the local protein structure with an average perturbation of 0.049 ppm in the conserved amphipathic region from residue Trp60 to Arg85 (three times that of the standard deviation threshold, σ_0_). For residue Leu73, directly adjacent to the site of lipidation, the CSP value reached 0.175 ppm (ten times the threshold value). Surprisingly, lipidation at Cys72 also seemed to have an effect on the disordered, soluble N-terminal domain, where 5 residues (Val5, Gln6, Phe9, Ser10, and Arg49) had a CSP value twice that of the standard deviation threshold. Although unexpected, this could be due in part to a change in how the disordered N-terminal region interacts with the DPC micelle when the amphipathic region of His_6_-IFITM3 is anchored to the membrane via a lipid analogue. This analysis indicates that although the most distinct change in the structure of IFITM3 is in the region proximal to the site of modification, lipidation at Cys72 can cause distal changes in the IFITM3 backbone. These results demonstrate that Cys72-lipidation can significantly affect the backbone conformation of IFITM3 and provides a potential structural link between the lipid modification and increased antiviral activity of Cys72-lipidated IFITM3.

### Biochemical analysis of IFITM3 membrane association

Beyond a structural understanding of how lipidation affects IFITM3, we explored how lipidation changed the interaction of IFITM3 with the local membrane environment. In order to directly test how lipidation affects the interaction of the amphipathic domain of IFITM3 with a phospholipid bilayer, we employed a classical flotation assay to measure the association of this region with reconstituted liposomes. In brief, IFITM3 protein samples were mixed with pre-formed liposomes and applied to a density gradient for ultracentrifugation. The liposomes, which contain low-density buffer, float to the top of the density gradient with any membrane-associated protein, whereas soluble protein remains at the bottom of the ultracentrifuge tube. The ultracentrifuged sample can be fractionated and visualized via SDS-PAGE to determine if the protein is associated with the liposomes (**Figure 5A**).

**Figure 5.**
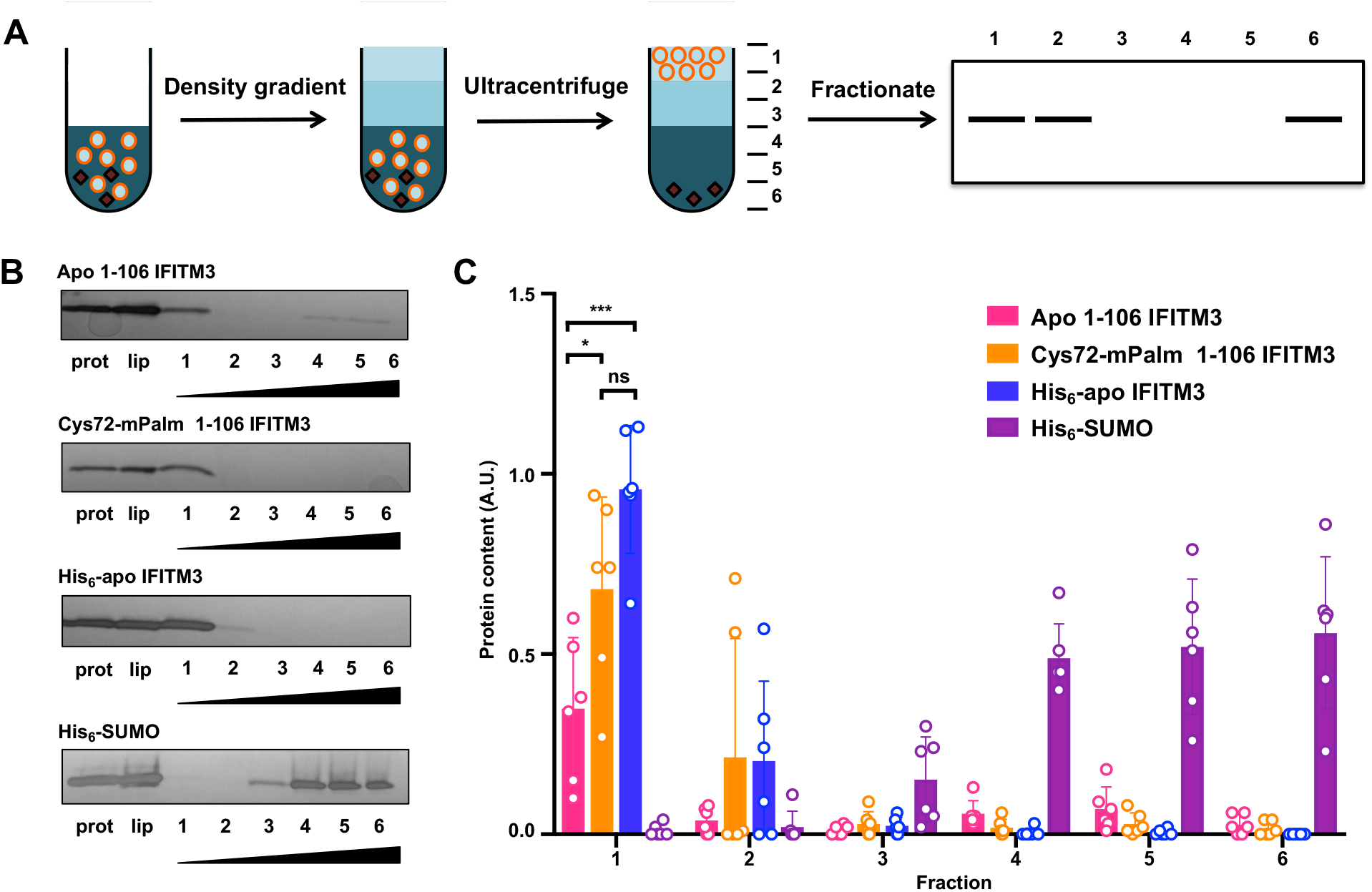
Flotation assay with lipidated variants of truncated IFITM3. A) Schematic of a liposome flotation assay. The liposome (orange circles) and protein (black) mixture in 40% Histodenz was first applied to the bottom of the ultracentrifuge tube. Then, layers of 30% Histodenz and buffer were stacked on top of the liposome mixture to form a density gradient. The samples were then ultracentrifuged and fractionated. Protein associated with the buoyant liposomes could be visualized in the top fraction, while soluble protein remained in the bottom fractions. B) Representative fractionation gels of apo 1-106 IFITM3, Cys72-mPalm 1-106 IFITM3, full length His_6_-apo IFITM3, and His_6_-SUMO flotation assay with lanes for protein alone, liposome control, and fractionated sample was visualized using silver stain. Wedges indicate increasing Histodenz concentration. C) Quantification of protein fractions from flotation assay normalized to the liposome loading control. The mean was obtained from six independent experiments (n=6) and is represented ± SD. Data were analyzed by one-way ANOVA with a post-hoc Tukey’s test (*p < 0.05, ***p < 0.0005). The following figure supplement is available for figure 5: **Figure supplement 1**. MALDI analysis of apo 1-106 IFITM3 and Cys72-mPalm 1-106 IFITM3 constructs.

To focus on the activity of the disordered and amphipathic regions of IFITM3, we designed a truncated construct (Met1-Leu106) that lacked the transmembrane domain and fused it to a cleavable His_6_-SUMO domain for improved expression. After purifying His_6_-SUMO-apo IFITM3 and His_6_-SUMO-Cys72 IFITM3 from the crude lysate with a Ni-NTA column, the SUMO domain was cleaved by ULP1 and IFITM3 was coupled to mPalm as previously described. The 1-106 IFITM3 constructs were then separated from the His_6_-SUMO domain by size exclusion. Successful purification and mPalm coupling was confirmed by MALDI analysis (**Figure 5 – figure supplement 2**). After purification, these truncated IFITM3 constructs were mixed with preformed 100 nm liposomes and dialyzed overnight to remove any detergent. Full length His_6_-apo IFITM3 and His_6_-SUMO were used as positive and negative controls, respectively. For the flotation assay, these protein-containing liposomes were added to a Histodenz density gradient and centrifuged at 150,000 x g for 3 hours. The sample was then fractionated and analyzed using SDS-PAGE to determine whether the protein was associated with the liposome. Since full length IFITM3 is an integral membrane protein, it fully associated with liposomes and was only found in the top fraction (**Figure 5B**). His_6_-SUMO, which has no membrane binding properties, was found in the bottom three fractions (**Figure 5B**). For the truncated IFITM3 constructs, liposome binding was correlated with lipidation at Cys72 (**Figure 5B**). Indeed, when we quantified the protein content across the density gradient, on average only 35% of apo 1-106 IFITM3 was present in the liposome-associated fraction, whereas Cys72-mPalm 1-106 IFITM3 was largely found in this first fraction (on average 68% of the total liposome protein content) (**Figure 5C**). These results show that lipidation at Cys72 increases the affinity of the amphipathic region of IFITM3 with the phospholipid bilayer, indicating that lipidation may be required to anchor the extended amphipathic region to the membrane, which is consistent with our molecular dynamics simulations (**Figure 2**).

### Site-specific chemical lipidation and antiviral activity of IFITM3 in mammalian cells

To evaluate site-specific lipidation of IFITM3 in mammalian cells, we employed genetic code expansion (J. W. Chin, 2014; Liu & Schultz, 2010) and bioorthogonal labeling (Lang & Chin, 2014; Prescher & Bertozzi, 2005) to install a stable *S*-palmitoylation mimic *in vivo* (**Figure 6A**). Since maleimide-palmitate cannot be used to site-specifically label IFITM3 in mammalian cells, we employed Inverse-Demand Diels Alder ligation (Selvaraj & Fox, 2013; Šečkutė & Devaraj, 2013) and tetrazine-lipid derivatives recently developed by our laboratories (**Li Y et al in review, Supplemental File**). For these studies, we used unnatural amino acids (UAAs) with strained alkenes or alkynes including axial trans-cyclooct-2-ene-lysine (2’-aTCOK) and exo-bicyclo[6.1.0]nonyne-lysine (exo-BCNK) as well as an alkyne-lysine control (AlkK) (**Figure 6B**). We mutated Cys72 of HA-tagged IFITM3 to the amber codon TAG to generate the construct HA-IFITM3-Cys72TAG. HEK293T cells were co-transfected with plasmids encoding an aminoacyl-tRNA synthetase/tRNA pair Mm-PylRS-AF (Y306A, Y384F)/Pyl-tRNA and HA-IFITM3-Cys72TAG in the absence or presence of the UAAs. Western blot analysis showed efficient expression of full length IFITM3 with the UAAs (**Figure 6 – figure supplement 1**). Following 16-hour transfection, cells were treated with a tetrazine-lipid mimic (Tz-6) (**Figure 6C**) for bioorthogonal tetrazine ligation reaction in live cells. The alkyne on Tz-6 enabled in-gel fluorescence profiling of chemically lipidated IFITM3 (**Figure 6D**). A Cu^I^-catalyzed azide alkyne cycloaddition (CuAAC) reaction of the cell lysate with azide-rhodamine enabled quantification of the efficiency of tetrazine ligation with different UAAs (**Figure 6A**,**D**). IFITM3 modified with an alkyne-containing UAA (AlkK) was used as a positive control for the CuAAC reaction and functioned as a benchmark to compare the efficiency of UAA incorporation and tetrazine ligation for the other UAA-containing constructs (**Figure 6E**). Both the UAAs showed successful chemical labeling of IFITM3, with the highest labeling efficiency for exo-BCNK (**Figure 6E**), consistent with our studies of H-Ras (**Li Y et al in review, Supplemental File**). We tested the chemical ligation efficiency at the other Cys sites of IFITM3, showing robust labeling of HA-IFITM3-Cys71TAG and HA-IFITM3-Cys105TAG with the tetrazine-lipid mimic Tz-6 (**Figure 6 – figure supplement 2**). We also evaluated the ligation efficiency with other tetrazine-lipid analogues of shorter aliphatic chain length (Tz-1) and higher hydrophilicity (Tz-PEG) (**Figure 6 – figure supplement 3a**). HA-IFITM3-Cys72TAG showed efficient chemical labeling independent of the nature of the tetrazine derivatives (**Figure 6 – figure supplement 3b**,**c**). These results show that the UAAs are successfully incorporated at specific sites in IFITM3 via genetic code expansion and can be modified with tetrazine derivatives for site-specific chemical lipidation of IFITM3 in live cells.

**Figure 6.**
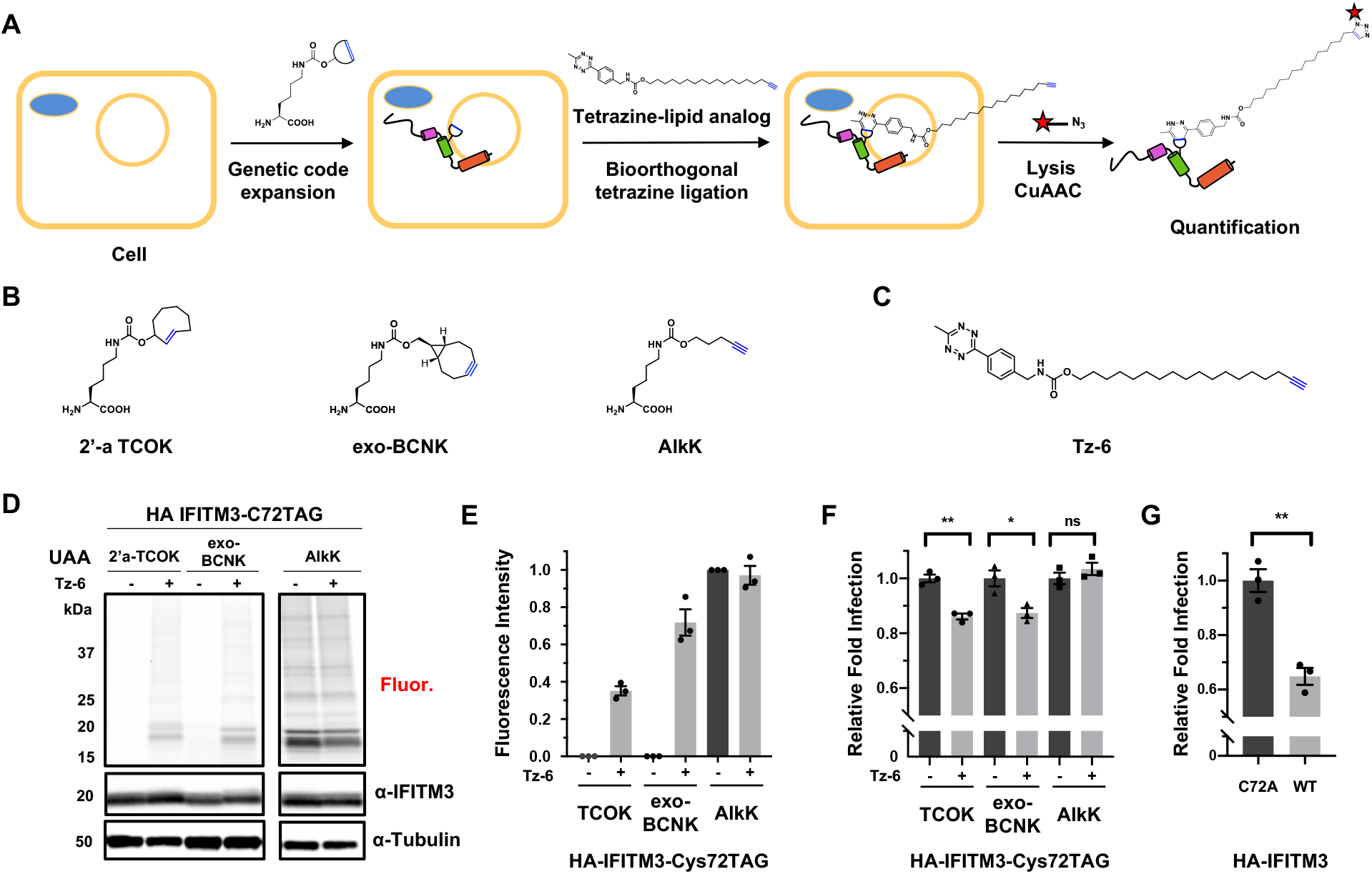
Chemical lipidation and anti-viral activity of IFITM3 in cells. A) Scheme for the site-specific lipidation of IFITM3 via genetic code expansion for unnatural amino acid incorporation and bioorthogonal tetrazine ligation reaction. B) Chemical structures of unnatural amino acids (UAAs) used for genetic code expansion and C) fatty acyl tetrazine analog Tz-6 used for tetrazine ligation to UAAs. D) In-gel fluorescence profiling of chemical lipidation tetrazine ligation efficiency in cells. HEK293T cells were transfected with plasmids encoding an aminoacyl-tRNA synthetase/tRNA pair Mm-PylRS-AF (Y306A, Y384F)/Pyl-tRNA and HA-IFITM3-Cys72TAG in the absence or presence of UAAs, after which cells were treated with 20 μM fatty acyl tetrazine Tz-6. The cell lysates were further reacted with azide-rhodamine for in-gel fluorescence profiling of chemically lipidated IFITM3. An anti-IFITM3 blot showed IFITM3 expression levels in each condition. An anti-tubulin western blot was used as a protein loading control. E) Quantification of efficiency of in-cell tetrazine ligation reaction. Data represents mean ± S.E.M. for three independent experiments. F) Quantification of influenza A virus (IAV) infection assay of HA-IFITM3 Cys72TAG-expressing cells with and without Tz-6 ligation for a panel of UAAs. Cells expressing HA-IFITM3-Cys72TAG with different UAAs were treated with fatty acyl tetrazine analog Tz-6, then infected with IAV. Virus nucleoprotein (NP) and HA-IFITM3 protein levels were examined by flow cytometry using anti-NP and anti-IFITM3 staining, respectively. Transfected cells expressing IFITM3 were gated and analyzed for percentage of infection. Relative fold infection for IFITM3 expressing cells was calculated with and without Tz-6-treated samples. Data represents mean ± S.E.M. for three independent experiments. Data were analyzed by unpaired Student’s t-test (ns p > 0.05, *p < 0.05, **p < 0.005). G) Quantification of IAV infection assay of HA-IFITM3 wild type (WT) and Cys72 to Ala (C72A) mutant HA-IFITM3 transfected cells. Cells expressing HA-IFITM3 were infected with IAV and analyzed by flow cytometry as previously described. Data represents mean ± S.E.M. for three independent experiments. Data were analyzed by unpaired Student’s t-test (**p < 0.005). The following figure supplements are available for figure 6: **Figure supplement 1**. Western blot analysis of HA-IFITM3-Cys72TAG expression with different unnatural amino acids (UAAs). **Figure supplement 2**. In-gel fluorescence profiling of tetrazine ligation efficiency at different Cys positions in IFITM3 in cells. **Figure supplement 3**. In-gel fluorescence profiling of tetrazine ligation efficiency with different tetrazine derivatives in cells. **Figure supplement 4**. Subcellular localization of chemically modified HA-IFITM3-Cys72TAG.

We next evaluated the virus susceptibility of HEK293T cells expressing lipidated IFITM3-Cys72TAG. Cells were infected by H1N1 influenza virus (IAV) for 6 hours, and viral infection was measured by staining of viral nucleoprotein (NP) and HA-IFITM3 by flow cytometry. IFITM3 with AlkK at Cys72 cannot undergo tetrazine ligation and showed no change in antiviral activity on addition of Tz-6 (**Figure 6F**). However, cells expressing IFITM3 constructs containing 2’-aTCOK and exo-BCNK showed decreased viral susceptibility upon treatment with Tz-6 (**Figure 6F**), restoring approximately 35% of the antiviral activity of wild type IFITM3 when compared with a Cys72 to Ala loss-of-function mutant (**Figure 6G**). Immunofluorescence imaging of the cells expressing IFITM3 with exo-BCNK at Cys72 show endolysosomal localization similar to endogenous IFITM3 in HeLa cells (**Figure 6 – figure supplement 4**). Moreover, labeling with tetrazine-lipid mimic Tz-6 showed no evident difference in subcellular localization (**Figure 6 – figure supplement 4**), suggesting the changes induced by site-specific lipidation may be on local membrane interactions rather than drastic changes in cellular distribution. Notably, this live cell chemical lipidation approach provides the first evidence for gain-of-function by site-specific lipidation of IFITM3 in mammalian cells and underscores the significance of Cys72 *S*-palmitoylation in IFITM3 antiviral activity.

## DISCUSSION

IFITMs are key immune effectors conserved across vertebrates that protect against a broad range of viral pathogens. *S*-palmitoylation was previously found to be essential for the antiviral activity of IFITM3, but the direct effects of palmitoylation on its structure and cellular activity have not been explored due to a lack of accessible chemical tools. Since S-palmitoylation is required for IFITM3 antiviral activity, a better understanding of site-specific lipidation is central to delineating its mechanism of action. By developing new methods for the generation of lipidated IFITM3 both *in vitro* and *in vivo*, we have been able to directly probe the biophysical and cellular consequences of IFITM3 *S-* palmitoylation.

The study of *S*-fatty acylated proteins *in vitro* and in cells has been challenging due to the hydrophobicity of the modification as well as its reversibility and substoichiometric levels in cells (Linder & Deschenes, 2007; Resh, 2006). To address these challenges, site-selective bioconjugation methods, protein semi-synthesis, and genetic code expansion have been employed to attach lipid analogs to proteins (Hang & Linder, 2011). Using maleimide-palmitate, we have been able to generate lipid-modified recombinant IFITM3 for a range of biochemical and biophysical studies. Molecular dynamics simulations of *S*-palmitoylated IFITM3 and maleimide-palmitate modified IFITM3 were conducted to determine if maleimide palmitate is an appropriate mimic for a native *S*-palmitoyl modification. Both the natural palmitoyl thioester and the synthetic mimic yielded similar structural changes in our computational models, indicating that maleimide-palmitate is a reasonable substitute for *S*-palmitoylation. Surprisingly, the molecular dynamics simulations also revealed a significant change in the structure of the amphipathic region of IFITM3 when lipidated, with the stabilization of the functionally significant AH1 region. Indeed, when Cys72-maleimide palmitate modified IFITM3 was compared with unmodified IFITM3 using NMR, we found structural changes both locally and in the disordered N-terminal region. Furthermore, a flotation assay revealed the disordered and amphipathic regions of Cys72-mPalm IFITM3 had increased association with the membrane bilayer. We note that maleimide-palmitate modified IFITM3 does not contain the native thioester linkage, which could be addressed through biophysical studies of site-specifically *S*-palmitoylated IFITM3 from new protein semisynthesis methods (Harmand, Pattabiraman, & Bode, 2017; D. L. Huang et al., 2020).

For cellular studies, protein semi-synthesis (Rocks et al., 2005; 2010) of site-specifically lipidated Ras isoforms has been instrumental in elucidating the reversible features of S-palmitoylated peripheral membrane proteins in mammalian cells. However, IFITMs are S-palmitoylated type IV membrane proteins that may not insert into cellular membranes properly and are therefore not ideal for microinjection studies. More recently, genetic code expansion has been used to site-specifically functionalize proteins with lipid mimics (**Li Y et al in review, Supplemental File**). Here, we have leveraged the method developed by Li Y *et al*. to chemically lipidate cellular IFITM3, expanding the scope of the technique to integral membrane proteins. This has allowed us to conduct the first antiviral gain-of-function assay of lipidated IFITM3 in mammalian cells, confirming the importance of site-specific *S*-fatty acylation in the regulation of IFITM3 antiviral activity.

Together, our results provide the first direct evidence that lipidation can change the biophysical properties of IFITM3. As the disruption of AH1 is known to attenuate the antiviral activity of IFITM3 (Chesarino et al., 2017) and may aid in the remodeling of the endocytic membrane to prevent virus-endosome fusion (Guo et al., 2020), our work suggests that *S*-palmitoylation at Cys72 could enhance the antiviral activity of IFITM3 directly through the stabilization of AH1. Furthermore, lipidation of IFITM3 at Cys72 has been shown to be essential for the engagement of invading viral particles by IFITM3 (Spence et al., 2019; Suddala et al., 2019). AH1 stabilization by Cys72 palmitoylation may provide a structural mechanism for IFITM3 subcellular partitioning during infection. As our current *in vitro* work focuses on IFITM3 in detergent and simplified lipid environments, in the future we can use our novel maleimide-palmitate modified constructs in more physiologically relevant lipid environments to measure subdomain localization and lipidation dependent effects on the biophysical properties of the membrane. Furthermore, live cell imaging studies could be conducted with our specifically lipidated constructs by using dual unnatural amino acid incorporation for both lipid modification and fluorescent labeling of IFITM3. Given the unique role of IFITMs in SARS-CoV-2 infection (Prelli Bozzo et al., 2020; Shi et al., 2020; X. Zhang et al., 2020a), understanding the regulation of IFITM3 via *S*-palmitoylation may be important for determining the host immune response to COVID-19. As *S*-palmitoylated IFITM3 is also a key viral restriction factor in common zoonotic disease vectors such as bats (Benfield et al., 2020), a mechanistic understanding of lipidated IFITM3 may also be important for the prevention of future disease outbreaks. In summary, our findings illuminate important links between the protein structure, membrane association, and antiviral activity of IFITM3, and underscore the significance of integrated *in silico*, chemical, biophysical, and cellular studies of *S*-fatty acylation.

## MATERIALS AND METHODS

### Molecular dynamics simulations

All Molecular dynamics simulation systems were prepared using the CHARMM36 force field for protein and lipid (B. R. Brooks et al., 2009; Klauda et al., 2010; Venable et al., 2014). The initial Apo IFITM3 structure was built using IFITM3 sequence data (Ling et al., 2016) and typical α-helical ϕ/ψ angles (−57.8° for ϕ and -47.0° for ψ) for AH1, AH2, and TM helices. The initial structure was then equilibrated in an implicit solvent (GBSW) environment with CHARMM (B. R. Brooks et al., 2009; Im, Feig, & Brooks, 2003a; Im, Lee, & Brooks, 2003b). The positional restraints were applied during the equilibration to place TM in membrane region and AH1 and AH2 on the membrane surface. After the equilibration, three simulation systems (Apo, Cys72-Palm, and Cys72-mPalm) were prepared using CHARMM-GUI (Jo, Kim, Iyer, & Im, 2008) *Membrane Builder* (Jo, Lim, Klauda, & Im, 2009; Emilia L Wu et al., 2014). The palmitoyl group was added at Cys72 during the *Membrane Builder* system building process for Cys72-Palm system. The maleimide group in maleimide-palmitate was parameterized by analogy of C3/C5 dyes in CHARMM-GUI (Jo et al., 2014) and added to the IFITM3 structure for Cys72-mPalm system. TIP3P water (Jorgensen, Chandrasekhar, Madura, Impey, & Klein, 1983) and 150 mM KCl ion were added to the bulk region with for the ionic strength of cellular environment. The upper and lower leaflet contains 94 number of DMPC lipids with the box size of 80 × 80 x 100 Å^3^. Equilibration of system were conducted with CHARMM-GUI standard protocol, and the productions were simulated with OpenMM 7.1 (Eastman et al., 2017). The particle-mesh Ewald method (Essmann et al., 1995) was used for long-range electrostatic interaction, and the SHAKE algorithm (Ryckaert, Ciccotti, & Berendsen, 1977) was utilized for fixing all bonds including hydrogen atom. The temperature of each systems was held at 308.15 K using Langevin dynamics (Goga, Rzepiela, de Vries, Marrink, & Berendsen, 2012), and pressure was stayed at 1 bar under the semi-isotropic Monte-Carlo barostat method (Åqvist, Wennerström, Nervall, Bjelic, & Brandsdal, 2004; Chow & Ferguson, 1995) with a 5 ps^-1^ coupling frequency. A force-based switching method (Steinbach & Brooks, 1994) was used for the van der Waals interactions under 10 to 12 angstrom cut-off range. To reduce the uncertainty of sequence-based structure, 1.5 µs of preliminary simulation was performed and the coordinate information from that of simulation was used for the later simulation. Among 1 µs of production runs, the last 500 ns trajectories were used for analysis, and each system has 3 replicas to obtain better sampling.

### Maleimide palmitate synthesis

A round bottom flask was charged with 0.68 g of triphenylphosphine (2.57 mmol, 0.9 eq.) and 17.5 ml of tetrahydrofuran. The flask was placed under argon and cooled to -78 °C. 1.18 ml of the 40% solution of diethyl azodicarboxylate in toluene (2.57 mmol, 0.9 eq.) were added over a period of 3 minutes. The resulting mixture was stirred for 5 minutes, after which a solution of 0.7 g of hexadecan-1-ol (2.87 mmol, 1 eq.) in a minimal amount of THF (prepared in an argon purged vial) was added over a period of 1 minute. The resulting solution was stirred for 5 minutes. The flask’s septum was then removed under the protection of an argon curtain and 0.125 g of neopentyl alcohol (1.43 mmol, 0.5 eq.) and 0.25 g of maleimide (2.57 mmol, 0.9 eq.) were added as solids. The flask was closed again under argon and the reaction mixture was stirred for 5 minutes, after which the cooling bath was removed and the reaction stirred at room temperature for 16 hours, then at 40 °C for 2 hours. After full conversion was indicated by TLC, the solvent was evaporated in vacuo. The resulting solid was purified by silica flash chromatography (loading and elution in dichloromethane). N-1-hexadecylmaleimide was obtained after high vaccum drying (0.481 g, 52%). H NMR (400 MHz, CDCl3) δ ppm 0.88 (t, J=6.59 Hz, 13 3H) 1.25 (br.s.,26H) 1.53-1.60 (m,2H) 3.51 (t,J=7.23Hz,2H) 6.68 (s,2H). CNMR (100MHz, CDCl3): δ ppm 14.09, 22.67, 26.73, 28.52, 29.11, 29.34, 29.46, 29.53, 29.64, 31.90, 37.93, 134.00, 170.87. This experimental procedure was adapted from Matuszak *et al*. (Matuszak, Muccioli, Labar, & Lambert, 2009).

### IFITM3 purification and maleimide-palmitate coupling

To express His_6_-IFITM3, the *E. coli* codon optimized gene was inserted into a pET-28c vector and transformed into BL21(DE3) *E. coli*, after which cells were grown in LB to 0.8 OD_600_ before being induced with 1 mM IPTG overnight at 18°C. Cells were then lysed with a probe sonicator in 2% TX-100, 25 mM HEPES pH 7.4, 150 mM KCl and 1 mM TCEP. After ultracentrifugation to remove insoluble cell debris (30K, 30 min, 4°C), the lysate was diluted to <1% TX-100 with buffer and incubated with cobalt beads for 30 minutes at 4°C. After washing beads with 10x CV of 0.8% TX-100, 25 mM HEPES pH 7.4, 150 mM KCl, and 1 mM TCEP (Buffer A) and 10x CV Buffer A with 40 mM imidazole, the protein was eluted with 3x CV of Buffer B (Buffer A with 400 mM Imidazole). The sample was then transferred to a 15°C shaker, where it was incubated with 0.5 mM maleimide palmitate for 2 hours with a final concentration of 10% DMSO. Post incubation, the sample was diluted with Buffer A until the imidazole concentration dropped below 20 mM and was incubated with cobalt beads for another 30 minutes. Beads were washed with 10x CV DPC buffer (0.5% DPC, 25 mM HEPES pH 7.4, 150 mM KCl) before being eluted with DPC buffer + 400 mM imidazole.

### MALDI analysis

1 μL of the sample was mixed with 9 μL of matrix consisting of a saturated solution of α-cyano-4-hydroxycinnamic acid (4-HCCA) in a 1:3:2 (v/v/v) mixture of formic acid/water/isopropanol (FWI). An aliquot of 0.5 – 1 μL of this protein-matrix solution was spotted onto a MALDI plate precoated with an ultrathin layer of 4-HCCA matrix^84,85^. The sample spots were then washed for a few seconds with 2 μL of cold 0.1% aqueous trifluoroacetic acid (TFA) solution. MALDI spectra were acquired in linear, delayed extraction mode using a Spiral TOF JMS-S3000 (JEOL, Tokyo, Japan). The instrument is equipped with a Nd:YLF laser, delivering 10-Hz pulses at 349 nm. Delayed extraction time was set at 0.75 – 1 μs and acquisition was performed with a sampling rate of 2 ns. Each MALDI spectrum corresponded to an average of 1000 scans. Mass calibration was performed using horse myoglobin as protein calibrant with a technique of pseudo-internal calibration wherein a few laser shots on a calibrant spot near a sample spot were collected and averaged with the sample shots into a single spectrum. The spectra were processed and analyzed using MoverZ (Proteometrics, LLC).

### NMR sample preparation

To express His_6_-IFITM3, the *E. coli* codon optimized gene was inserted into a pET-28c vector and transformed into BL21(DE3) *E. coli*. For NMR experiments, cells were adapted to 100% D2O ^13^C^15^N M9 minimal media, before being induced with 1 mM IPTG overnight at 18°C. After induction, IFITM3 was purified and chemically lipidated as described above. After purification, samples were concentrated and dialyzed overnight into the final NMR buffer (0.5% DPC, 25 mM HEPES pH 7, 150 mM KCl). His_6_-apo IFITM3 spectra were recorded on 150 μM, 350 μM, and 1 mM samples. His_6_-Cys72-mPalm IFITM3 spectra were recorded on a sample at 350 μM.

### NMR spectroscopy and analysis

The NMR data was acquired at 35°C (308K) on Bruker 800 and 900 MHz AVANCE spectrometers equipped with TCI CryoProbes. The backbone resonances of ^2^H^13^C^15^N labeled IFITM3 with and without lipidation were assigned using a suite of multidimensional N^15^-TROSY based experiments: ^15^N-HSQC, HNCA, HNCO, HNCACB, HN(CO)CA, HN(CO)CACB, HNCACO and ^15^N-edited NOESY-HSQC. Spectra analysis was performed in CARA. The assignments were confirmed by recording 2D N^15^-TROSY on selectively N^15^-labeled samples (Ala, Ile, Val, Leu and Phe) with and without lipidation. The ^15^N, ^13^Cα, and ^13^C’ chemical shift assignments were then used to predict the dihedral angles of the protein backbone using TALOS+ Shen et al., 2009). In the His_6_-apo IFITM3 sample, 70 out of 115 predicted dihedral angles were found to be “good” by TALOS+. Of the other residues, 19 residues were predicted to be dynamic (84% of which were in the disordered N-terminal region from residue Met1 to Val59) and 26 residues were found to be ambiguous. Out of those residues, 8 were in the structured C-terminal domain (Trp60 to Gly133) while 18 were in the disordered N-terminal region (Met1 to Val59). In the His_6_-Cys72-mPalm IFITM3 sample, 68 out of 106 predicted dihedral angles were found to be “good” by TALOS+, while 25 residues were predicted to be dynamic (21 residues in the disordered N-terminal region, 4 residues in the structured C-terminal region). 13 residues were found to be ambiguous, with 5 in the structured C-terminal domain and 8 in the disordered N-terminal region. For the chemical shift perturbation analysis, the chemical shift differences were calculated using a weighted average of the Euclidean distance moved:

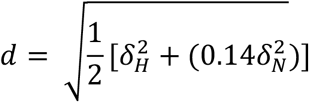

The standard deviation of the change in chemical shift across the protein was calculated and used as a threshold value (σ_0_) to identify residues with significant changes in chemical shift (Williamson, 2013).

### Truncated construct purification

Truncated IFITM3 constructs were inserted into a SUMO containing pET28c(+) plasmid (gift from the Lima lab at MSKCC). Lysis and His-tag purification was completed in the same manner as the full length constructs. After His purification, the samples were incubated with ULP1 (1:250) overnight at 4°C. The next morning, the samples were incubated at 15°C for 2 hours with maleimide palmitate. After coupling, the samples were concentrated and buffer exchanged into 1% OG, 25 mM HEPES, and 150 mM KCl over size exclusion (SEC200).

### Flotation assay

Liposomes were prepared by extruding 5mM lipid stock (80:20 POPC:Cholesterol) through a 100nm filter. As maleimide-palmitate modified 1-106 IFITM3 was unstable without detergent, the liposomes were mixed 1:1 with protein in 1% OG, 25 mM HEPES, 150 mM KCl so the final protein concentration was 1 μM and the detergent saturated but did not solubilize the liposomes (Rigaud & Lévy, 2003). The liposomes were then incubated at RT for 1 hour to promote protein incorporation, after which the liposomes were dialyzed into 25 mM HEPES and 150 mM KCl overnight at 4°C. Buffer was exchanged after 1 hour, then again for 2 hours the next morning. For the flotation assay, liposomes were mixed with 80% histodenz solution so the final concentration was 40%. 600 μL of this 40% histodenz solution was layered in the bottom of the ultracentrifuge tube, followed by 400 μL 30% histodenz and 200 μL buffer. The samples were ultracentrifuged at 150,000 x g for 3 hours at 4°C, after which each sample was fractionated and acetone precipitated overnight at -20°C. The protein pellets were resuspended in SDS and quantified using silver stain.

### Cell culture and reagents

HEK293T cells were purchased from ATCC and cultured in Dulbecco’s Modified Eagle’s Medium (DMEM, high glucose; Gibco) supplemented with 10% fetal bovine serum (FBS; VWR). Influenza A/PR/8/34 (H1N1) (10100374) was from Charles River Laboratories. Influenza A virus nucleoprotein antibody [AA5H] (ab20343) was from Abcam. Alexa-Fluor 647 Antibody Labeling Kit (A20186) was ordered from Life Technologies. Anti-HA antibody conjugated to Alexa-Fluor 594 was purchased from Life Technologies. Anti-HA HRP-conjugated antibody was ordered from Roche.

### Plasmid transfection

pCMV-Mm-PylRS-WT plasmid was kindly provided by Professor Peng R. Chen at Peking University (Zhang et al., 2011). pCMV-FLAG-Mm-PylRS-AF (Y306A and Y384F double mutant of wild-type PylRS) were generated in the lab by site-directed mutagenesis of pCMV-Mm-PylRS-WT and introducing an in-frame N-terminal FLAG tag. Human IFITM3 cDNA was purchased from Open Biosystems and PCR cloned into pCMV-HA vector (Clontech). All mutants of IFITM3 were generated by using QuikChange II Site-Directed Mutagenesis Kit (Agilent Technologies, #200523). Lipofectamine 3000 from Thermo Scientific was used for transfection of HEK293T cells.

### Chemical lipidation of IFITM3

HEK293T cells were seeded on 6-well plates and cultured overnight. The next day cells were co-transfected with the plasmid encoding aminoacyl-tRNA synthetase/tRNA pair Mm-PylRS-AF (Y306A, Y384F)/Pyl-tRNA (0.5 µg) and HA-IFITM3-Cys72TAG (0.5 µg) using 3 µL Lipofectamine 3000 in complete cell growth media containing UAAs (100 μM) for 16 h. Then cells were treated with tetrazine-lipid (20 μM) for 2 h. Cells were lysed with 4% SDS lysis buffer (4% SDS, 150 mM NaCl, 50 mM triethanolamine pH 7.4, Roche protease inhibitor, benzonase). Protein concentrations were determined by the BCA assay (Pierce). Protein concentration of cell lysate was normalized to 1.6 μg/ml.

### Fluorophore labeling of chemically lipidated IFITM3

45 μl of the cell lysate was treated with 5 μl of CuAAC reactant solution (0.5 μl of 10 mM azido-rhodamine (final concentration 100 μM), 1 μl of 50 mM freshly prepared CuSO_4_·5H_2_O in H_2_O (final concentration 1 mM, Sigma), 1 μl of 50 mM freshly prepared TCEP (final concentration 1 mM) and 2.5 μl of 10 mM Tris[(1-benzyl-1H-1,2,3-triazol-4-yl)methyl]amine (TBTA) (final concentration 500 μM)) was added. The samples were rocked at room temperature for 1 h.

### In-gel fluorescence profiling and western blot

Protein pellet was precipitated by using a mixture of methanol-chloroform-H_2_O (4:1.5:3, relative to sample volume). After mixing by inversion several times, samples were centrifuged at 20,000x g for 5 min at 4°C. Two separate phases were observed with a protein pellet between the two. After carefully removing the aqueous (top) layer, 1 mL of prechilled methanol was added and centrifuged. After another wash with methanol, the protein pellet was dried using speed-vacuum for 10 min. The pellet was resuspended in 1X Laemmli sample buffer (20 μl) and was heated for 10 min at 95 °C and separated by gel electrophoresis. In-gel fluorescence scanning was performed using a Bio-Rad ChemiDoc MP Imaging System. Gels were transferred to nitrocellulose membranes using BioRad Trans-Blot Semi-Dry Cell (20 V, 40 min), which were blocked with PBST (0.05% Tween-20 in PBS) containing 5% nonfat milk for 1 h at room temperature. The membranes were then incubated with anti-HA HRP conjugate (3F10, Roche, 1:2000 dilution) overnight at 4 °C. After overnight incubation, membranes were washed with PBST three times and developed using Bio-Rad Clarity Western ECL substrate and imaged with a Bio-Rad ChemiDoc MP Imager. Quantification of band intensities in fluorescence gels and Western blotting were performed with Image Lab (Bio-Rad). Data from three biological replicates were quantified and averaged for plotting.

### Influenza A virus infection assay

For the infection assay, HEK293T cells were transfected and treated as described above to express chemically lipidated IFITM3. Then the cells were infected with influenza virus A/PR/8/34 virus (H1N1) with MOI of 2.5. After 6 h, cells were washed with PBS, trypsinized, and collected in cluster tubes. Cells were washed again with PBS and then fixed with 240 µl of 4% PFA in PBS for 5 min. The fixed cells were permeabilized with 200 µl of 0.2% saponin in PBS for 10 min and then blocked with 200 µL of 0.2% BSA and 0.2% saponin in PBS for 10 min. Cells were treated with anti-influenza NP antibody conjugated to AlexaFluor-647 (1:250) and anti-HA antibody conjugated to AlexaFluor-594 (1:250) in 0.02% saponin in PBS. After three washes with 0.02% saponin in PBS, cells were resuspended in 100 µL PBS containing 0.2% BSA and 0.02% saponin. The samples were analyzed by flow cytometry (BD LSRII). Data analysis was performed using FlowJo software. All samples were first gated by IFITM3-positive staining, indicating successful transfection, and then NP-positive percentage of NP-positive staining, indicated successful infection.

## ACKNOWLEDGEMENTS

E. H. Garst and T. Das were supported by the Tri-Institutional Chemical Biology program through the NIH Chemistry-Biology Training Grant T32 GM115327. A. Percher was supported by an NSF Graduate Research Fellowship. We thank P.D.R Olinares and B.T. Chait for technical assistance and discussion of MALDI analysis. We thank M. Griffin, J. Yount, and T. Walz for helpful discussion and input into this manuscript. The data collected at NYSBC was made possible by a grant from ORIP/NIH facility improvement grant CO6RR015495. The 900 MHz NMR spectrometers were purchased with funds from NIH grant P41GM066354 and the New York State Assembly. T. Peng acknowledges support from National Natural Science Foundation of China (21778010) and Shenzhen Science and Technology Innovation Committee (JCYJ20170412150832022). This work was supported by NSF MCB-1810695 (to W. Im) and NIH-NIGMS R01GM087544 grant (to H.C. Hang).

## Notes

### Competing Interest Statement

The authors have declared no competing interest.

