## Supplementary Information for "Site-specific lipidation enhances IFITM3 membrane interactions and antiviral activity"


Emma Garst[1,2†], Hwayoung Lee[3†], Tandrila Das[1,2†], Shibani Bhattacharya[4], Avital Percher[1], Rafal Wiewiora[2,5], Isaac Witte[1], Yumeng Li[6], Michael Goger[4], Tao Peng[1,6], Wonpil Im[3], Howard C. Hang[1,7]*

[1] Laboratory of Chemical Biology and Microbial Pathogenesis, The Rockefeller University, New York, New York 10065, United States.

[2] Tri-Institutional Ph.D. Program in Chemical Biology, New York, NY 10065, United States.

[3] Department of Biological Sciences, Chemistry, and Bioengineering, Lehigh University, Bethlehem, PA 18015, United States.

[4] New York Structural Biology Center, New York, NY 10027, United States.

[5] Memorial Sloan Kettering Cancer Center, New York, NY 10065, United States.

[6] State Key Laboratory of Chemical Oncogenomics, School of Chemical Biology and Biotechnology, Peking University Shenzhen Graduate School, Shenzhen 518055, China.

[7] Departments of Immunology and Microbiology and Chemistry, Scripps Research, La Jolla, CA 92037, United States.

†These authors contributed equally.

### SUPPLEMENTARY FIGURES

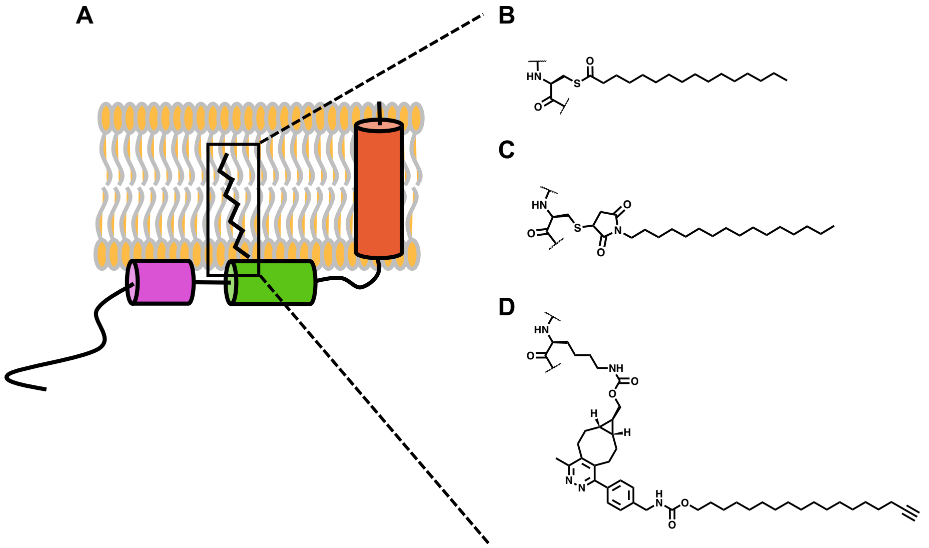


**Figure 1 – figure supplement 1. Structure and lipidation of IFITM3.** A) Cartoon of Cys72-*S*-palmitoylated IFITM3 based on previously published structural work. NMR and EPR analysis indicate IFITM3 is a type IV single-pass transmembrane protein with an extended amphipathic region from residue Trp60 to Arg85. B) Chemical structure of the *S*-palmitoyl post translational modification. In cells, IFITM3 is *S*-palmitoylated at Cys72. C) Chemical structure of cysteine conjugated maleimide palmitate (Cys72-mPalm-IFITM3). Cys72-mPalm-IFITM3 was used throughout our work for computational, structural, and biochemical analysis. D) Chemical structure of Cys72-lipidated IFITM3 modified through genetic code expansion and in cell bioorthogonal tetrazine ligation (Cys72-endo-BCNK-Tz-6 IFITM3). Cys72-endo-BCNK-Tz-6 IFITM3 was utilized for in cell antiviral gain-of-function assays.

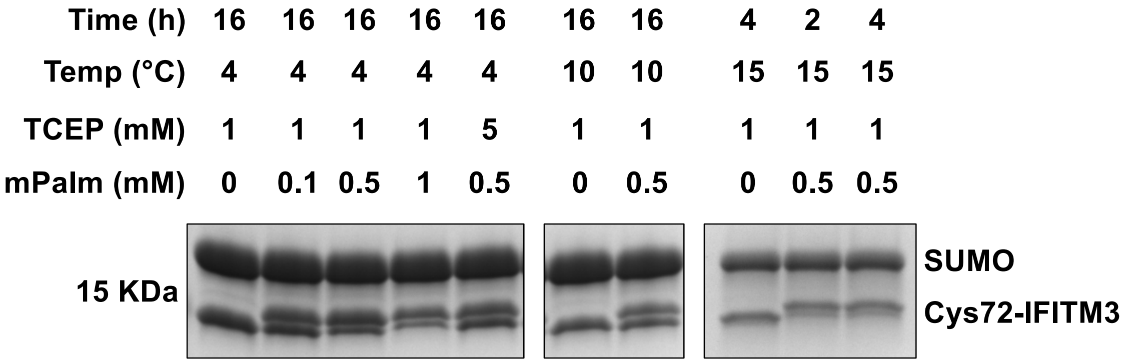


**Figure 3 – figure supplement 1. Optimization of Cys72-IFITM3 maleimide-palmitate coupling reaction.** A number of conditions were tested for the maleimide-palmitate (mPalm) coupling reaction, such as time, temperature, TCEP concentration, and mPalm concentration. Successful coupling was monitored by a mass shift on an SDS PAGE gel. At 4°C and 10°C, the maleimide coupling reaction did not go to completion after overnight coincubation with Cys72-IFITM3, as shown by a clear double band. Neither increasing the mPalm concentration nor increasing the TCEP concentration pushed the reaction to completion. However, at 15°C the reaction went to completion in 2 hours.

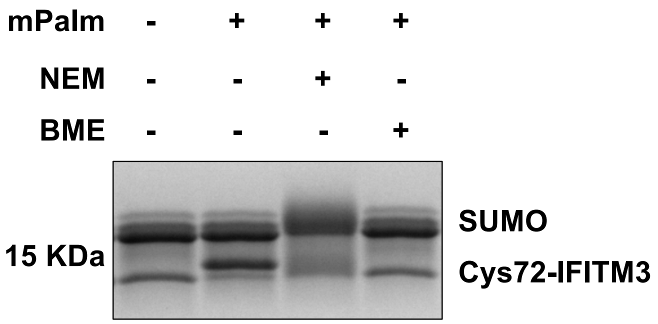


**Figure 3 – figure supplement 2. Maleimide-palmitate coupling with Cys72-IFITM3 can be blocked or quenched by *N*-ethyl maleimide (NEM) or β-mercaptoethanol (BME) respectively.** In brief, to test the ability of NEM and BME to block the mPalm-IFITM3 coupling reaction, Cys72-IFITM3 was pre-incubated with 20mM NEM for 30 minutes prior to mPalm coupling (NEM+ condition), or mPalm was preincubated with 5mM BME for 30 minutes before the Cys72-IFITM3 coupling reaction (BME+ condition). Maleimide-palmitate coupling to Cys72-IFITM3 was monitored via mass shift in an SDS PAGE gel. While a clear band shift was seen when Cys72-IFITM3 was coincubated directly with mPalm as previously described, coincubation with NEM or BME appeared to abrogate the effect.

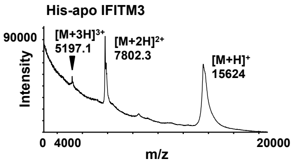


| | $[M+H]^+$ | $[M+2H]^{2+}$ | $[M+3H]^{3+}$ |
|---|---|---|---|
| **Expected mass (m/z)** | 15765 | 7883.5 | 5255.0 |
| **Observed mass (m/z)** | 15624 | 7802.3 | 5197.1 |
| **Difference (Da)** | -141 | -81.2 | -57.9 |

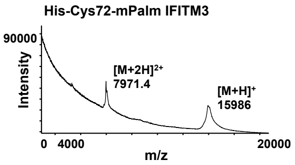


| | $[M+H]^+$ | $[M+2H]^{2+}$ | $[M+3H]^{3+}$ |
|---|---|---|---|
| **Expected mass (m/z)** | 16119 | 8060.3 | 5373.9 |
| **Observed mass (m/z)** | 15986 | 7971.4 | N/A |
| **Difference (Da)** | -133 | -88.9 | N/A |

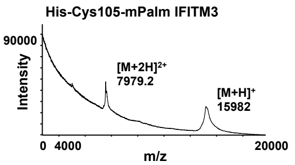


| | $[M+H]^+$ | $[M+2H]^{2+}$ | $[M+3H]^{3+}$ |
|---|---|---|---|
| **Expected mass (m/z)** | 16119 | 8060.3 | 5373.9 |
| **Observed mass (m/z)** | 15982 | 7979.2 | N/A |
| **Difference (Da)** | -137 | -81.1 | N/A |

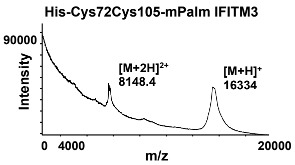


| | $[M+H]^+$ | $[M+2H]^{2+}$ | $[M+3H]^{3+}$ |
|---|---|---|---|
| **Expected mass (m/z)** | 16472 | 8237.1 | 5491.7 |
| **Observed mass (m/z)** | 16334 | 8148.4 | N/A |
| **Difference (Da)** | -138 | -88.7 | N/A |

**Figure 3 – figure supplement 3. MALDI quantification of $His_6$-apo IFITM3 and panel of $His_6$-IFITM3 lipidated variants.** The observed and expected masses of each $His_6$-IFITM3 variant were compared in order to confirm the efficacy of the mPalm coupling reaction. Comparison between spectra reveals a 362 Da increase in mass for $His_6$-Cys72-mPalm-IFITM3 and 358 Da increase in mass for $His_6$-Cys105-mPalm-IFITM3 when compared to unlipidated IFITM3 ($His_6$-apo-IFITM3). The dually lipidated IFITM3 construct ($His_6$-Cys72Cys105-mPalm-IFITM3) has a mass increase of 710 Da over the unmodified construct. Each construct had a net 130 – 180 Da loss in mass when compared to the expected molecular weight, indicating the loss of a terminal Met during expression or purification.

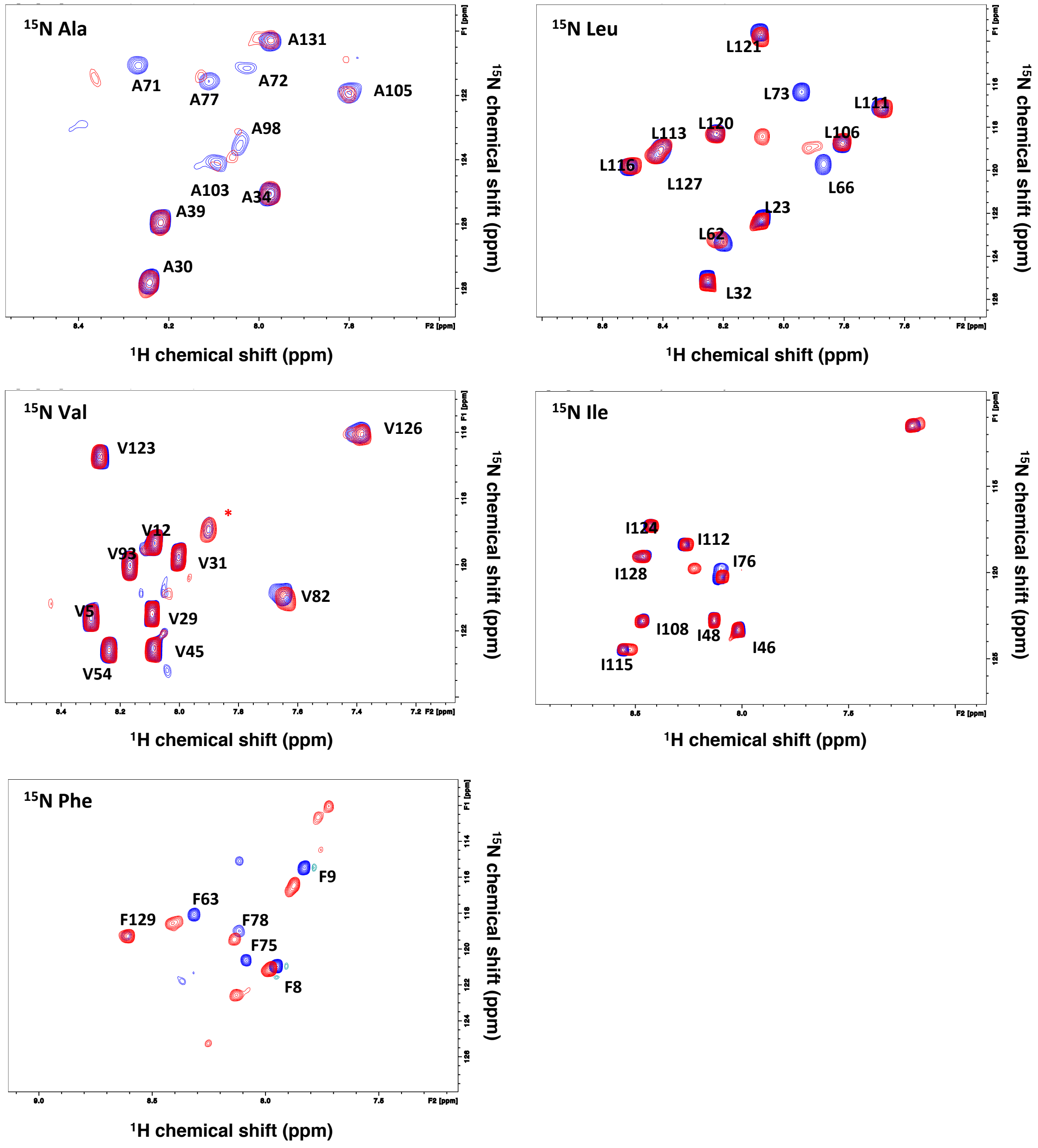


**Figure 4 – figure supplement 1. $^{15}$N specific labeling of His$_6$-apo and His$_6$-Cys-mPalm IFITM3 for backbone assignment.** $^1$H-$^{15}$N TROSY spectra were collected of $^{15}$N-specifically labeled His$_6$-apo (blue) and His$_6$-Cys72-mPalm (red) IFITM3 samples. The residues used in specific labeling were (clockwise from top left): $^{15}$N Ala, $^{15}$N Leu, $^{15}$N Ile, $^{15}$N Phe, and $^{15}$N Val.

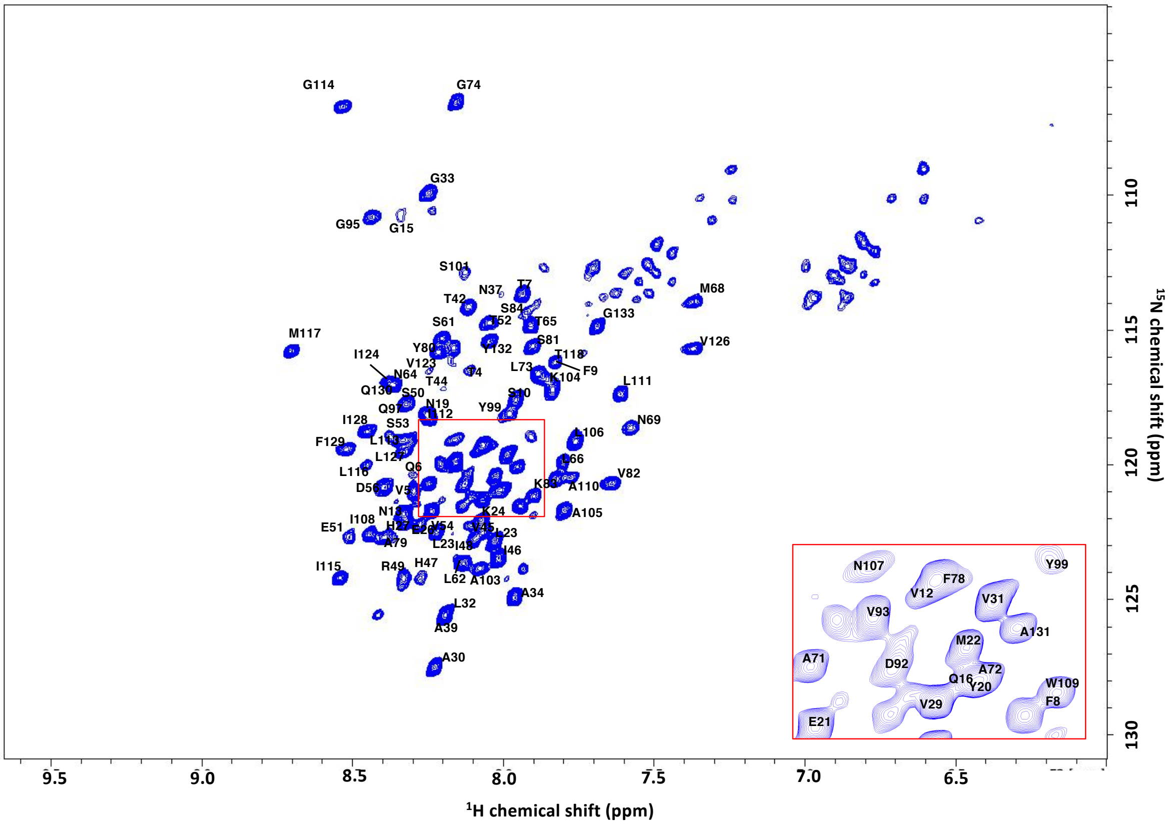


**Figure 4 – figure supplement 2. Backbone resonance assignment of His$_6$-apo IFITM3 overlayed on $^{1}$H-$^{15}$N TROSY spectra.** All His$_6$-apo IFITM3 assignment experiments were conducted on $^{2}$H$^{13}$N$^{15}$C labeled His$_6$-apo IFITM3 at 35ºC in 25 mM HEPES pH 7, 150 mM KCl, and 0.5% DPC. Residue assignment was achieved via the following 3D experiments: 15N-HSQC, HNCA, HNCO, HNCACB, HNCOCA, HNCOCACB, and HNCACO (see supplementary methods).

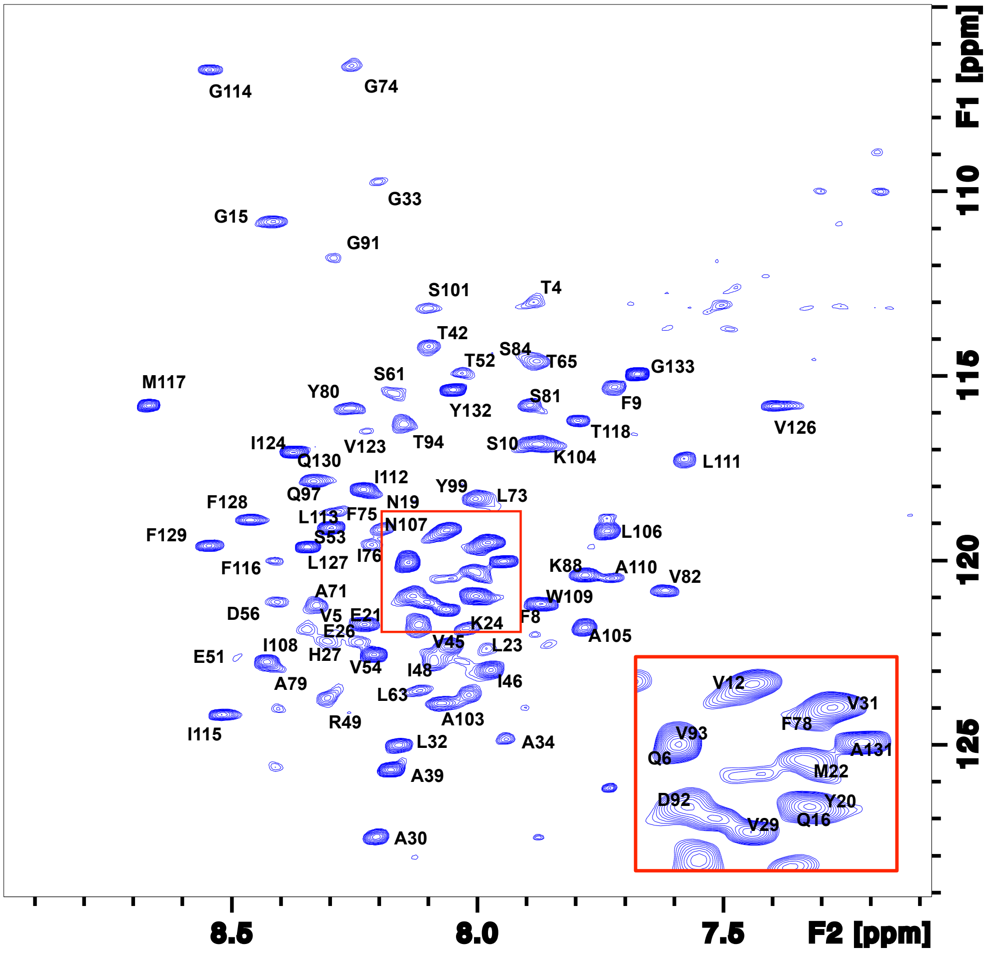


**Figure 4 – figure supplement 3. Backbone resonance assignment of His$_6$-Cys72-mPalm IFITM3 overlayed on $^1$H-$^{15}$N TROSY spectra.** All His$_6$-Cys72-mPalm IFITM3 assignment experiments were conducted on $^2$H$^{13}$N$^{15}$C labeled His$_6$-apo IFITM3 at 35ºC in 25 mM HEPES pH 7, 150 mM KCl, and 0.5% DPC. Residue assignment was achieved via the following 3D experiments: 15N-HSQC, HNCA, HNCO, HNCACB, HNCOCA, HNCOCACB, and HNCACO (see supplementary methods).

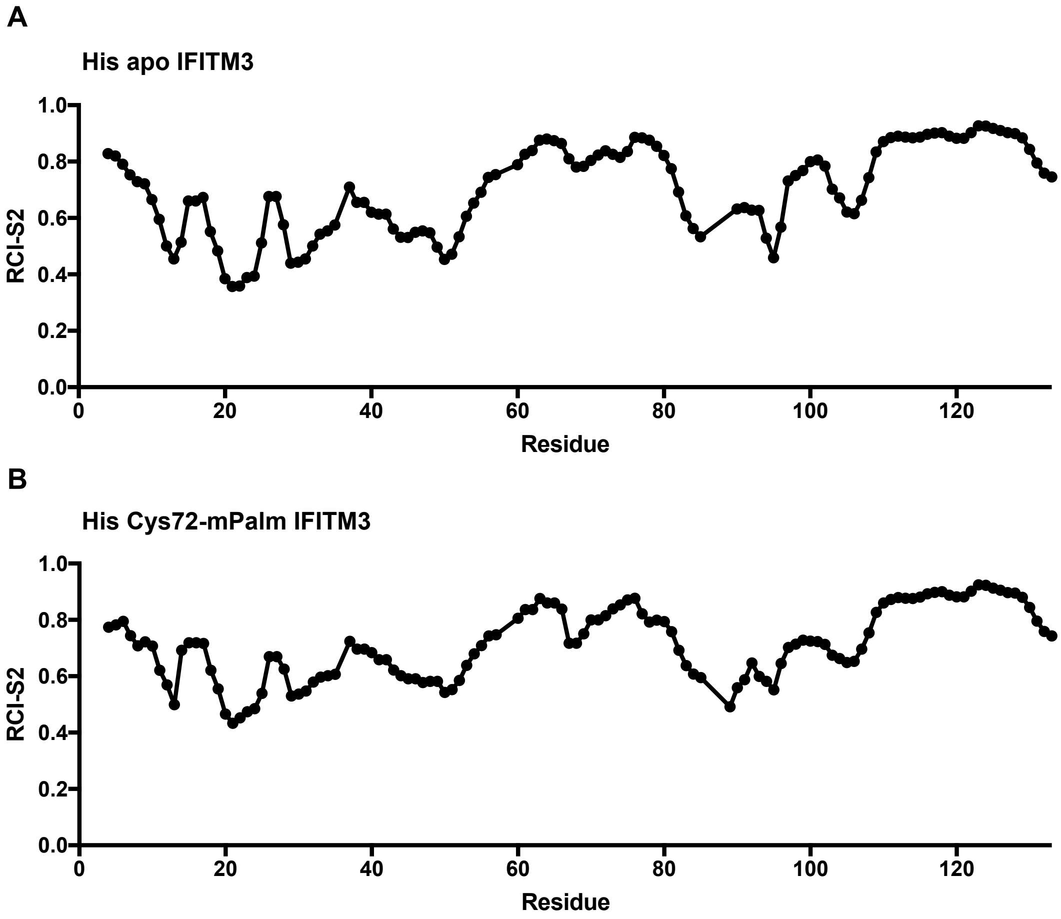


**Figure 4 – figure supplement 4. RCI analysis of His$_6$-apo IFITM3 and His$_6$-Cys72-mPalm IFITM3.** The random coil index predicted order parameter S2 was calculated from protein chemical shifts determined via backbone assignment of His$_6$-apo and His$_6$-Cys72-mPalm IFITM3. The calculation was preformed by the web-based TALOS+ analysis program[1] as described in Berjanskii *et al.*[2]

**Apo IFITM3**

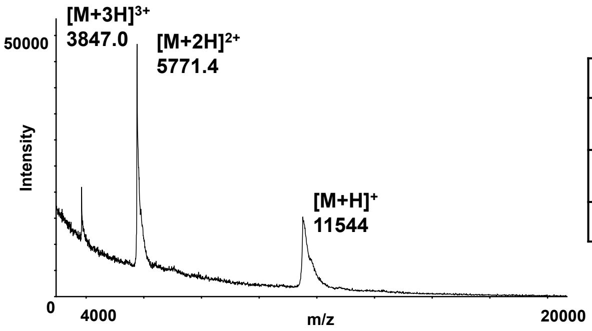


| | $[M+H]^+$ | $[M+2H]^{2+}$ | $[M+3H]^{3+}$ |
|---|---|---|---|
| **Expected mass (m/z)** | 11542 | 5772.1 | 3848.7 |
| **Observed mass (m/z)** | 11544 | 5771.4 | 3847.0 |
| **Difference (Da)** | +2 | -0.7 | -1.7 |

**Cys72-mPalm IFITM3**

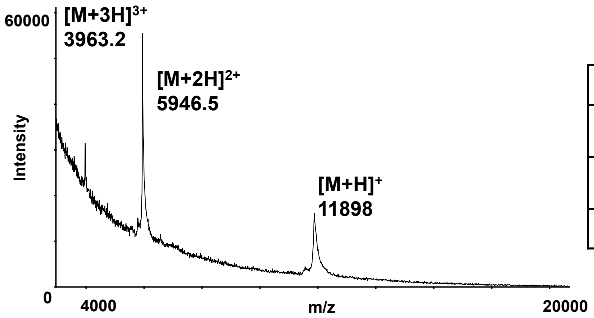


| | $[M+H]^+$ | $[M+2H]^{2+}$ | $[M+3H]^{3+}$ |
|---|---|---|---|
| **Expected mass (m/z)** | 11896 | 5948.9 | 3966.2 |
| **Observed mass (m/z)** | 11898 | 5946.5 | 3963.2 |
| **Difference (Da)** | +2 | -2.4 | -3 |

**Figure 5 – figure supplement 1. MALDI analysis of apo 1-106 IFITM3 and Cys72-mPalm 1-106 IFITM3.** MALDI spectra of apo 1-106 IFITM3 and Cys72-mPalm 1-106 IFITM3 were collected in linear, delayed extraction mode a 2 ns sampling rate and 0.75-1 $\mu$s delay. Samples were calibrated internally with a horse myoglobin protein standard. The observed and expected masses of the apo 1-106 and Cys72-mPalm 1-106 IFITM3 variants were compared in order to confirm the efficacy of the mPalm coupling reaction. Comparison between spectra revealed a 354 Da increase in mass, indicating the addition of one mPalm group. The expected and observed masses for both constructs were in accordance, with less than a ± 3 Da difference for each peak.

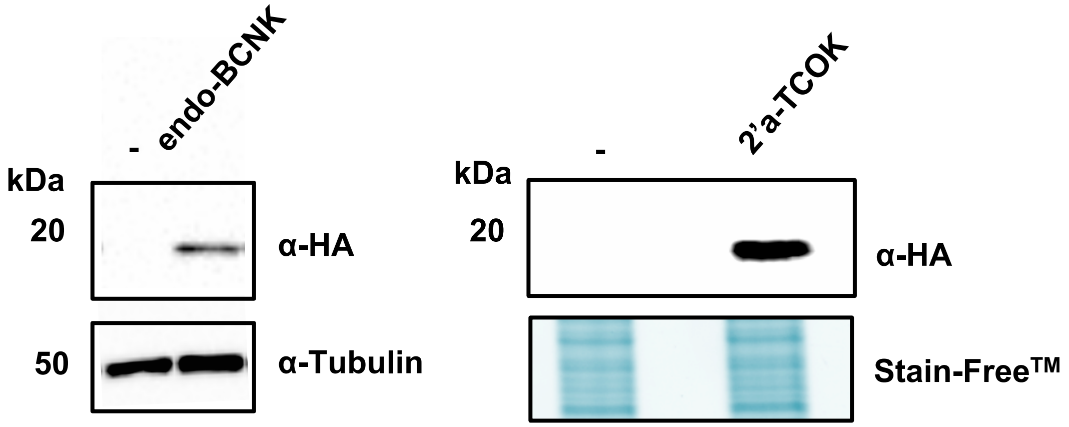


**Figure 6 – figure supplement 1. Western blot analysis of HA-IFITM3-Cys72TAG expression with different unnatural amino acids (UAAs).** HEK293T cells were transfected with plasmids encoding an aminoacyl-tRNA synthetase/tRNA pair Mm-PylRS-AF (Y306A, Y384F)/Pyl-tRNA and HA-IFITM3-Cys72TAG in the absence or presence of different UAAs. Anti-HA blot shows efficient genetic code expansion for HA-IFITM3-C72TAG expression with different UAAs. Anti-Tubulin blot or Stain-Free$^{TM}$ gel (Bio-Rad) is shown as protein loading control.

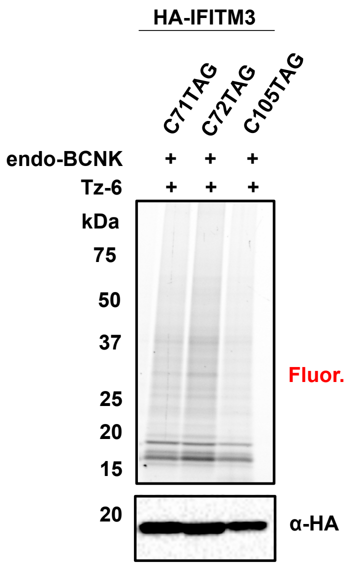


**Figure 6 – figure supplement 2. In-gel fluorescence profiling of tetrazine ligation efficiency at different Cys positions in IFITM3 in cells.** HEK293T cells were transfected with plasmids encoding an aminoacyl-tRNA synthetase/tRNA pair Mm-PylRS-AF (Y306A, Y384F)/Pyl-tRNA and HA-IFITM3-Cys71TAG or Cys72TAG or Cys105TAG in the presence of endo-BCNK, after which cells were treated with fatty acyl tetrazine Tz-6. The cell lysates were further reacted with azide-rhodamine for in-gel fluorescence profiling of chemically lipidated IFITM3. An anti-HA blot revealed similar IFITM3 expression levels in each condition.

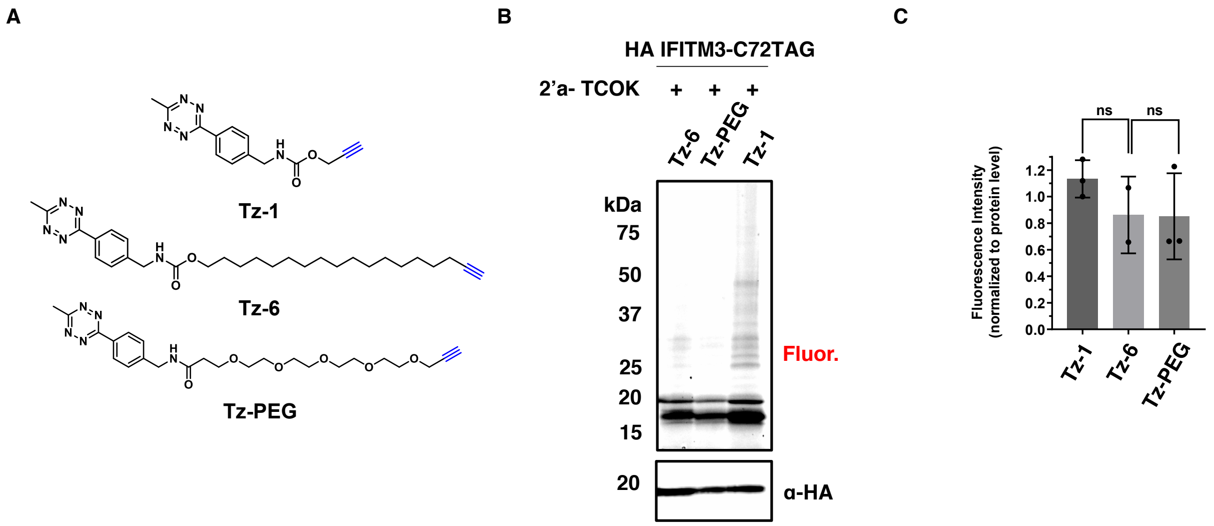


**Figure 6 – figure supplement 3. In-gel fluorescence profiling of tetrazine ligation efficiency with different tetrazine derivatives in cells.** A) Structures of the different tetrazine derivatives used for tetrazine ligation to UAAs. B) HEK293T cells were transfected with plasmids encoding an aminoacyl-tRNA synthetase/tRNA pair Mm-PylRS-AF (Y306A, Y384F)/Pyl-tRNA and HA-IFITM3-Cys72TAG in the presence of 2'a-TCOK, after which cells were treated with different tetrazine analogs. The cell lysates were further reacted with azide-rhodamine for in-gel fluorescence profiling of chemically modified IFITM3. Anti-HA blot shows IFITM3 expression level in each condition. C) Quantification of efficiency of in-cell tetrazine ligation reaction. Data represents mean ± S.E.M. for three independent experiments. Data were analyzed by unpaired Student's t-test (ns $p > 0.05$).

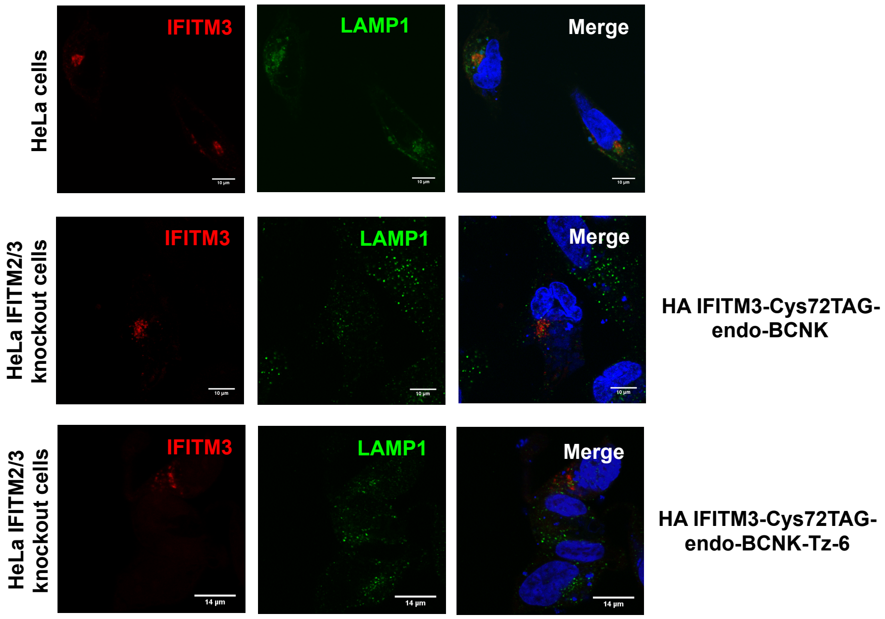


**Figure 6 – figure supplement 4. Subcellular localization of chemically modified HA-IFITM3-Cys72TAG.** IFITM 2/3 KO HeLa cells were transfected with plasmids encoding an aminoacyl-tRNA synthetase/tRNA pair Mm-PylRS-AF (Y306A, Y384F)/Pyl-tRNA and HA-IFITM3-Cys72TAG in the presence of endo-BCNK, after which cells were treated with tetrazine analogue Tz-6. Cells were stained with anti-IFITM3 antibody and anti-LAMP1 antibody and DAPI for nucleus. Top row, localization of endogenous IFITM3 (red) and LAMP1 (green) in HeLa cells; middle row, LAMP1 (green) and HA-IFITM3 with endo-BCNK at Cys72 (red) expressed in HeLa IFITM2/3 knockout cells; bottom row, LAMP1 (green) and HA-IFITM3 chemically lipidated with Tz-6 in HeLa IFITM2/3 knockout cells.
